# CD3^+^ T-cell: CD14^+^monocyte complexes are dynamic and increased with HIV and glucose intolerance

**DOI:** 10.1101/2023.04.24.538020

**Authors:** Laventa M. Obare, Joshua Simmons, Jared Oakes, Xiuqi Zhang, Cindy Nochowicz, Stephen Priest, Samuel S. Bailin, Christopher M. Warren, Mona Mashayekhi, Heather K. Beasley, Jianqiang Shao, Leslie M. Meenderink, Quanhu Sheng, Joey Stolze, Rama Gangula, Tarek Absi, Yan Ru Su, Kit Neikirk, Abha Chopra, Curtis L. Gabriel, Tecla Temu, Suman Pakala, Erin M. Wilfong, Sara Gianella, Elizabeth J. Phillips, David G. Harrison, Antentor Hinton, Spyros A. Kalams, Annet Kirabo, Simon A. Mallal, John R. Koethe, Celestine N. Wanjalla

## Abstract

An increased risk of cardiometabolic disease accompanies persistent systemic inflammation. Yet, the innate and adaptive immune system features in persons who develop these conditions remain poorly defined. Doublets, or cell-cell complexes, are routinely eliminated from flow cytometric and other immune phenotyping analyses, which limits our understanding of their relationship to disease states. Using well-characterized clinical cohorts, including participants with controlled HIV as a model for chronic inflammation and increased immune cell interactions, we show that circulating CD14^+^ monocytes complexed to CD3^+^ T cells are dynamic, biologically relevant, and increased in individuals with diabetes after adjusting for confounding factors. The complexes form functional immune synapses with increased expression of proinflammatory cytokines and greater glucose utilization. Furthermore, in persons with HIV, the CD3^+^T-cell: CD14^+^monocyte complexes had more HIV copies compared to matched CD14^+^ monocytes or CD4^+^ T cells alone. Our results demonstrate that circulating CD3^+^T-cell:CD14^+^monocyte pairs represent dynamic cellular interactions that may contribute to inflammation and cardiometabolic disease pathogenesis and may originate or be maintained, in part, by chronic viral infections. These findings provide a foundation for future studies investigating mechanisms linking T cell-monocyte cell-cell complexes to developing immune-mediated diseases, including HIV and diabetes.

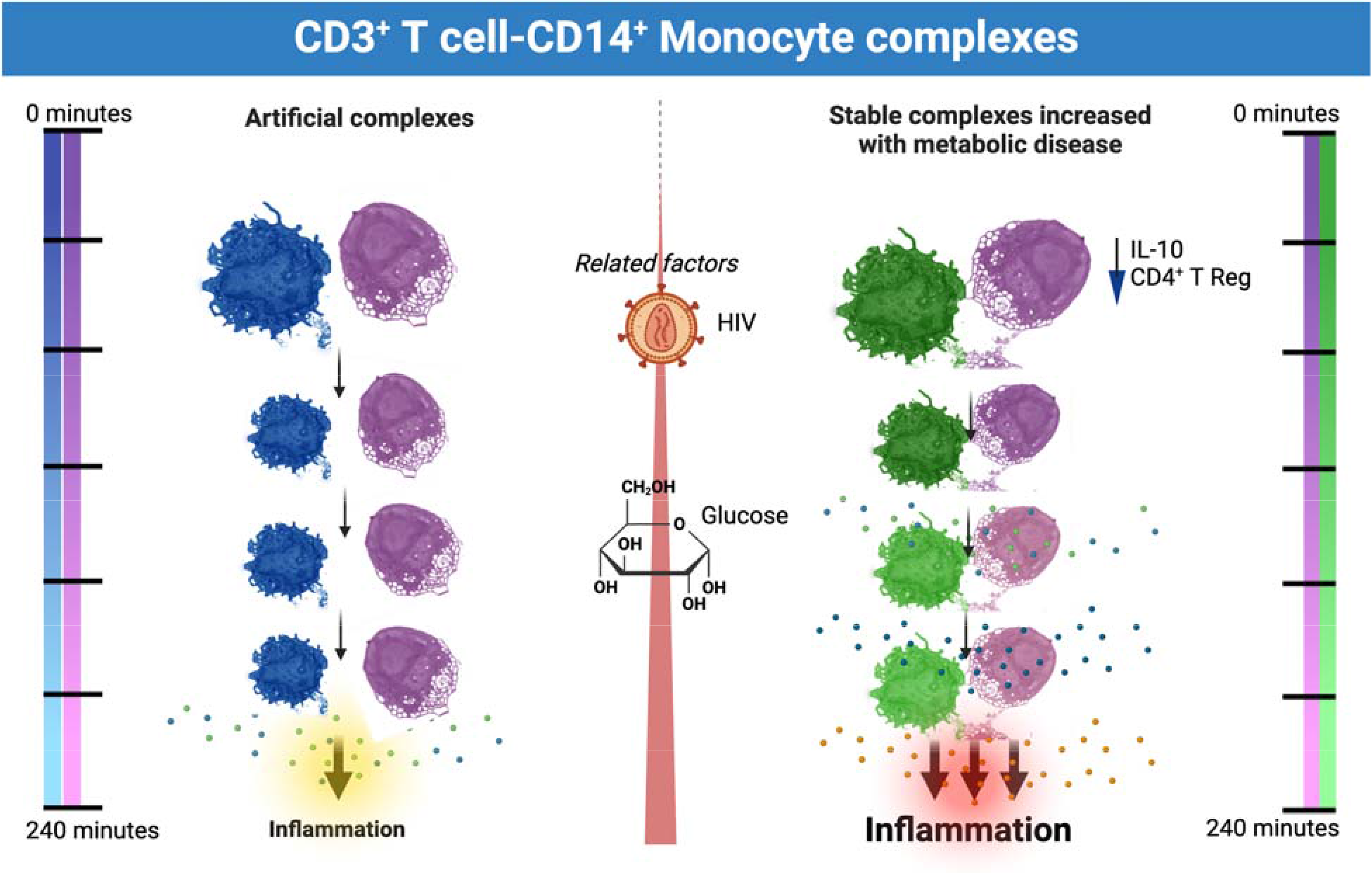

**Highlights:**

- Circulating CD3^+^ CD14^+^ T cell-monocyte complexes are higher in individuals with diabetes.
- CD3^+^ CD14^+^ T cell-monocytes complexes comprise a heterogenous group of functional and dynamic cell-cell interactions.
- The proportion of CD3^+^ CD14^+^ T cell-monocyte complexes is positively associated with fasting blood glucose and negatively with plasma IL-10 levels and CD4^+^ T regulatory cells.
- CD3^+^ CD14^+^ T cell-monocyte complexes are metabolically flexible and can utilize both glycolysis and oxidative phosphorylation for their energy requirements.
- In persons with treated HIV, CD3^+^ CD14^+^ T cell-monocytes have more detectable HIV DNA than circulating CD4^+^ T cells alone.

## Introduction

Chronic inflammation is linked to diseases like diabetes and atherosclerosis, which can worsen in people living with HIV (PLWH) due to elevated cytokine and chemokine levels despite antiretroviral therapy (ART) (Alcaide et al., 2013; Bailin et al., 2022; Bailin et al., 2020; Grome et al., 2017; Kundu et al., 2022; Temu et al., 2020; Temu et al., 2021; Wanjalla, Mashayekhi, et al., 2021). Traditional analysis of immune cells using flow cytometry often misses interacting cell populations, dismissing cell aggregates as artifacts of sample processing (Hsue & Waters, 2019). However, recent evidence suggests the presence of significant immunologically relevant cell-cell complexes in various disease states, including tuberculosis and after vaccination with immunogenic vaccines such as the yellow fever vaccine (Burel et al., 2019; Gil-Manso et al., 2021) (Sivakumar et al., 2021). Importantly, these cell-cell complexes are not artifactual from cryopreservation of peripheral blood mononuclear cells (PBMCs), with a strong correlation between these complexes from fresh and cryopreserved samples (Burel et al., 2019). However, none of these studies have shown an association between cell-cell complexes and metabolic disease. Our study investigates these complexes in the context of controlled HIV, using it as a model to understand chronic immune activation’s effects on diseases like diabetes and cardiovascular disease.

## Methods

### Study Participants

All PLWH included in this study were previously recruited for the HIV, Adipose Tissue Immunology, *and Metabolism cohort* at the Vanderbilt Comprehensive Care clinic. The cohort has individuals without diabetes (hemoglobin A1c [HbA1c] <5.7% and fasting blood glucose [FBG] <100 mg/dl), with prediabetes (HbA1c 5.7-6.4% and/or FBG 100-125 mg/dl), and diabetes (HbA1c ≥6.5%, FBG≥126 mg/dl and/or on medications to treat diabetes). All PLWH were on ART with sustained viral suppression for at least 12 months before the study, with a CD4^+^ T cell count > 350 cells/ml (**Table S1**). The cohort excluded individuals with inflammatory illnesses, substance abuse, greater than 11 alcoholic drinks per week, and active hepatitis B/C. The study is registered at ClinicalTrials.gov (NCT04451980) (Wanjalla, Mashayekhi, et al., 2021). The second cohort has ten adults without HIV who were enrolled in an ongoing study to understand the role of immune cells in aging and cardiovascular disease (CVD) (**Table S2**). All studies were approved by the Vanderbilt University of Medicine Institutional Review Boards. Participants provided written informed consent. The investigators carried out studies using the United States Department of Health and Human Services guidelines.

### Sex as a biological variable

Our study examined males and females and similar parameters and measured and reported for both sexes.

### Mass cytometry

Mass cytometry was conducted on cryopreserved PBMCs using a validated 37-marker antibody panel (**Table S3**). PBMCs were stained for live/dead cells with Cisplatin, surface markers with a master mix, and fixed in paraformaldehyde (PFA). Post-fixation, cells were stored in methanol at −20°C, later stained with intracellular markers for 20 minutes at room temperature, followed by the addition of 2ul (250nM) of DNA intercalator (Ir) in phosphate-buffered saline (PBS) with 1.6% PFA. Just before running and analyzing the samples on the mass cytometer, we washed the cells in PBS, followed by Millipore water. For analysis, we resuspended 500,000 cells/ml of Millipore water. 1/10 volume of equilibration beads were added to the cells, which we then filtered and analyzed on the Helios. FCS files from the Helios cytometer were bead-normalized using the premessa R package’s normalizer GUI method (Gherardini, 2022). FCS files were analyzed in Flowjo to clean the data of debris (DNA-), Fluidigm beads (175++165++), and dead cells (cisplatin^+^). Data gating was performed using Flowjo (**Figure S1**), followed by downstream analysis in R programming language (version 4.2.1) and the flowCore package (Ellis B, 2022). We downsampled all samples and processed them through the following workflow: Subset parameters were transformed using the function asinh(x/5). A nearest neighbor search produced a weighted adjacency matrix with several nearest neighbors set to the dimension of subset + 1. (Arya S, 2019) The Leiden community detection algorithm was used to cluster the adjacency matrix. (Kelly, 2020) Uniform Manifold Approximation and Projection (UMAP) was done for subset visualization using the uwot R package (J, 2022).

### Tracking Responders Expanding populations (T-REX) and Marker Enrichment Modeling (MEM) of enriched features

The T-REX algorithm was performed as published (Barone et al., 2021). In brief, we classified cells of interest measured using mass cytometry including CD45^+^ cells, CD3^+^ CD4^+^ T cells, CD3^+^ CD8^+^ T cells, CD3^-^ CD19^-^ HLA-DR^+^ monocytes, and CD3^-^ CD56^+^ CD16^+^ NK cells. UMAP analyses were performed for concatenated non-diabetic participants (group 1) and prediabetic/diabetic participants (group 2). This was followed by K nearest neighbor (KNN) analyses to search for the nearest neighbors for each cell. The difference in the percent change per cell between group 1 and group 2 is calculated based on the abundance of these cells in each group in the KNN region. ≤5% and ≥95% changes in cell percentages were considered significant, which we clustered using Density-Based Spatial Clustering of Applications with Noise (DBSCAN). The phenotype of the clusters that were significantly different between the groups was determined using the MEM package.

### Flow cytometry

PBMCs were stained with fluorescently tagged antibodies as previously published (Wanjalla et al., 2019). In brief, thawed and washed PBMCs were stained with Aqua (Live/Dead marker) for 10 minutes at room temperature, followed by the addition of a mastermix of fluorescently tagged antibodies (**Table S2, BD Aria**). Stained cells were analyzed using a four-laser BD FACS ARIA III with a cell sorter. The two-dimensional gates used to sort CD3^+^T-cell: CD14^+^monocyte complexes are shown (**Figure S2A, supplemental Table S3**). The stained PBMCs were resuspended in PBS with RNA*later* to stabilize the RNA (ThermoFisher #AM7022). Single-cell indexed sorting was done using a 100μM nozzle to sort CD3^+^T-cell: CD14^+^monocyte complexes as a single entity into each 96 well plate as previously published (Wanjalla, McDonnell, et al., 2021b). For bulk sequencing, we performed a 4-way sort through a 100μM nozzle into four 1.5ml Eppendorf tubes (CD4^+^T-cells, CD8^+^T-cells, CD14^+^ monocytes and CD3^+^T-cells: CD14^+^monocyte complexes).

### Single Cell ENergetIc metabolism by profiling Translation inhibition (SCENITH) Assay

PBMCs were prepared for SCENITH as published (Argüello et al., 2020). We added the 10μl of inhibitors (oligomycin [1.5μM], 2DG [100mM], 2DG + Oligomycin) to the cells. Media only was included as a control. All samples were then incubated at 37°C for 30 minutes. We then added puromycin (10μM) to each condition (PBMCs with inhibitors) and incubated the PBMCs at 37°C for 45 minutes. After this, we washed the PBMCs in PBS, stained them with surface antibodies against CX3CR1 and CCR7, and stained them at 37°C for 15 minutes. The cells were then incubated with the master mix containing the other surface markers (**Cytek antibodies, Table S2**) for 20 minutes at room temperature. The cells were fixed with 4% PFA for 15 minutes at room temperature. We added 0.1% triton permeabilization solution and incubated the cells for 15 minutes. Anti-puromycin in permeabilization buffer was added to the cells for 15 minutes at room temperature. Cells were then washed and resuspended in PBS for analysis with Cytek Aurora.

### Droplet digital PCR

We sorted CD3^+^ CD4^+^ T cells, CD14^+^ monocytes, and CD3^+^ CD14^+^ T cell-monocyte complexes into separate Eppendorf tubes with PBS. Cells were pelleted and resuspended in lysis buffer [Tritonx100 (0.1%), Tris HCL (10mM), and Proteinase K (400ug/ml)] at 55°C for 10 hours. Additional proteinase K was added during the heat inactivation stage at 95°C for 5 minutes. For HIV DNA quantitation, we used LTR primers (forward primer -LTR 5’-AGC ACT CAA GGC AAG CTT TA-3’, and reverse primer -LTR 5’-TGT ACT GGG TCT CTC TGG TTA G-3’, and probe 5’-FAM-GCA GTG GGT TCC CTA GTT AGC CAG AGA G-3IABkFQ-3’) (Abana et al., 2017). HIV transcripts were quantified as copies/million cells. 19μl of the ddPCR SuperMix (LTR primers & RPP30 housekeeping gene primers and probes), and 6ul of cell lysates were mixed and aliquoted per well (96-well twin tec plate) and droplets generated with an AutoDG. Droplets were read using a plate reader, and the positive droplet threshold was manually set using the negative droplet control (media only).

### Time-lapse imaging

CD3^+^ CD14^+^ T cell-monocyte complexes were sorted as above and resuspended in RPMI with 10% fetal bovine serum (FBS). The cells were then plated on poly-L-Lysine pre-coated coverslips at a density of 15,000-40,000 complexes per 100μl media. The cells on the coverslip were placed in a 24-well plate, and time-lapse imaging was captured using an EVOS M5000 imaging system. Image J Version 1.53t 24 August 2022 was used for image analysis.

### Single-cell T-cell receptor (TCR) sequencing

Single-cell TCR sequencing involved sorting CD3^+^ CD14^+^ T cell-monocyte complexes, storing them at −80°C, and using uniquely tagged primers for reverse transcription (Wanjalla, McDonnell, et al., 2021a). cDNA amplification was performed with KAPA HiFi HotStart ReadyMix (Roche, Basel, Switzerland) (Grün, Kester, & van Oudenaarden, 2014; Islam et al., 2014; Kivioja et al., 2011). TCR gene expression was quantified via UMIs and nested PCRs targeting TCRαβ genes. After pooling and purifying the products, indexed sequencing libraries were created using Truseq adapters and quantified with the Jetseq qPCR Library Quantification Kit (Meridian Biosciences Inc., OH, USA). Samples were sequenced on an Illumina MiSeq with paired-end reads, quality-filtered, and demultiplexed. Reads were assigned to TCRA and TCRB loci and TCR clonotypes using MIXCR software (Bolotin et al., 2015), with data visualization by VGAS (Hertzman et al., 2021).

### Transmission electron microscopy (TEM)

CD3^+^ CD14^+^ T cell-monocyte complexes, CD3^+^ T cells, and CD14^+^ monocytes were sorted as above. For day 3 samples, we added RPMI media supplemented with human IL-2 [10ng/mL]. The cells were plated on a poly-L-lysine coated coverslip for 1-2 hours for doublet imaging. When the cells were bound to the coverslip, the media was aspirated, and then the cells were fixed with 2.5% glutaraldehyde solution in 0.1 M sodium cacodylate buffer (Neikirk et al., 2023). After secondary fixation, samples were washed for five minutes with 0.1 M sodium cacodylate buffer (7.3 pH). Followed by two five-minute washes with diH_2_O. While keeping all solutions and plates at room temperature, the samples were incubated with 2.5% uranyl acetate, diluted with H_2_O, at 4 °C overnight. The samples were dehydrated using an ethanol gradient series. After dehydration, the ethanol was replaced with Eponate 12™ mixed in 100% ethanol in a 1:1 solution, then incubated at room temperature for 30 mins. This was repeated three times for 1 hour using 100% Eponate 12^™^. The plates were finally placed in new media and cured in an oven at 70 °C overnight. The plates were cracked upon hardening, and the cells were separated by submerging the plate in liquid nitrogen. An 80 nm thickness jeweler’s saw was used to cut the block to fit in a Leica UC6 ultramicrotome sample holder. The section was placed on formvar-coated copper grids counterstained in 2% uranyl acetate for 2 mins. Then, the grids were counterstained by Reynold’s lead citrate for two minutes. TEM acquired images on either a JEOL JEM-1230, operating at 120 kV, or a JEOL 1400, operating at 80 kV (Lam et al., 2021).

### Single-cell RNA (scRNA) sequencing

PBMCs were thawed, washed with PBS, and incubated with Fc receptor-blocking solution. Surface antibody staining was performed (CD3 clone UCHT1 #300479, CD4 clone SK3 #344651, CD8a clone SK1 #344753, CD14 clone 63D3 #367137, CD16 clone 3G8 #302065, CD69 clone FN50 #310951), followed by encapsulation and barcoding using the Chromium Single Cell 5’ assay. Library preparation, cDNA amplification, and sequencing were done, aligning reads to the human genome. Cell identification and downstream analyses, including feature selection, PCA, and UMAP, were executed using Seurat V4 (Hao et al., 2021). Cells with abnormal gene or mitochondrial counts were filtered out. DoubletFinder identified doublets for exclusion. Differential gene expression analysis was conducted on sorted cell populations using WebGestalt (WEB-based Gene SeT Analysis Toolkit) (Wang, Duncan, Shi, & Zhang, 2013).

### Statistical analysis

This cross-sectional study establishes whether T cell-monocyte complexes are associated with metabolic disease variables and outcomes in PLWH. We reported summary statistics of clinical demographic characteristics using medians and interquartile ranges. Wilcoxon test was used to examine differences for continuous variables, and Pearson chi-squared test was used for categorical variables. We selected partial Spearman correlation for analysis because it is less sensitive to outliers. We used a nonparametric test, partial Spearman’s correlation analysis, to test the relationship between T cell-monocyte complexes and clinical variables, including hemoglobin A1C, fasting blood glucose, high-density lipoprotein (HDL), low-density lipoprotein (LDL), triglycerides, coronary arterial calcium, and fat volume (pericardial, subcutaneous, and visceral). We adjusted for possible confounders that could influence the relationship between the immune complexes and the outcomes. These included age, sex, and body mass index (BMI). We selected partial Spearman correlation because it is less sensitive to outliers. Similarly, we used partial Spearman’s correlation analysis to test the relationship between T cell-monocyte complexes and plasma cytokines. We adjusted for possible HIV-related confounders that could influence the immune complexes’ relationship with the plasma cytokines. These included CD4:CD8 ratio, hemoglobin A1C, and duration of years on ART.

Other statistical analyses comparing two continuous variables were performed using the Mann-Whitney U and Kruskal-Wallis tests, where more than two variables were compared. Statistical analysis in this study was performed in Graph Pad Prism version 9.5.0 and R version 4.2.1. Details of transcriptomic analysis above under single-cell sequencing.

### Data and code availability

Gene expression data from this study have been deposited in the NIH Gene Expression Omnibus (GEO) accession numbers: GSE229707 and GSE230276. Requests for further details on protocols and data included in this study are available upon request from the lead contact.

## Results

### Characteristics of PLWH

The HIV cohort comprised 38 individuals on ART with long-term suppression of plasma viremia: 14 without diabetes and 24 with prediabetes or diabetes (**Table S1**). Details of the cohort and clinical visit procedures were previously published (Wanjalla et al., 2019). In downstream analysis, PLWH with prediabetes and diabetes were combined into a single metabolic disease group. The characteristics of the groups were largely similar, except for parameters linked to glucose intolerance that we have highlighted and adjusted for in downstream analysis. These include body mass index (p<0.05), waist and hip circumference (p<0.05, p=0.01 respectively), and fasting blood sugar (p<0.001). Glucose-tolerant individuals were younger by about 10 years of age (p=0.1). There were no notable differences observed in HIV-related laboratory values (CD4 at ART start, CD4 at T cell enrollment, current ART, duration on ART, and hepatitis C antibody status. Cell-associated DNA and RNA were higher in PLWH without diabetes but not statistically significant (p=0.1). Similarly, visceral fat volume was higher with glucose intolerance but insignificant (p=0.1). Lastly, 33% of PLWH with diabetes had coronary arterial calcium (CAC), while none of the participants without HIV had CAC (p=0.02). The differences between PLWH with and without glucose intolerance included known risk factors associated with metabolic disease, including age, BMI, hip/waist circumference, fasting blood glucose, and CAC prevalence.

### Circulating cells of the innate and adaptive immune system differ by metabolic health

Mass cytometry examined immune cells in cryopreserved PBMCs of all PLWH (**Figure S1A**). We identified six primary clusters, including CD4^+^ T cells, senescent/cytotoxic CD4^+^ T cells (Wanjalla et al., 2019), CD8^+^ T cells, monocytes, B cells, and NK cells (**Figure 1A**). A comparison of clusters (abundance/size differences) between PLWH with diabetes/prediabetes, and those without revealed several clusters in participants with glucose intolerance that were fewer in PLWH without diabetes depicted with the red dotted circles, all p<0.05 (**Figure 1B, Figure S1B**). Other cell types that were more abundant in PLWH without diabetes included classical monocytes and CD14^+^ CD16^+/-^ Monocytes (**Table S4**). The heatmap in Figure 1C represents the median relative expression of immune markers on clusters and the median fold difference in cluster sizes. Magenta clusters were more abundant in prediabetic/diabetic PLWH, whereas CD4 T regulatory cell cluster marked with a blue oval was more abundant in non-diabetic PLWH. CGC^+^ CD4^+^ T cells, a population we have previously reported as associated with metabolic and cardiovascular disease conditions in controlled HIV, were also increased with diabetes (Wanjalla, Mashayekhi, et al., 2021; Wanjalla et al., 2019; Wanjalla, McDonnell, et al., 2021b). As previously published, we calculated the association constant between T cells/B cells and monocytes for the cell-cell complex clusters (Burel et al., 2019). To calculate the constant, we divided the proportion of the cell-cell complex by the proportion of T cells multiplied by the proportion of monocytes as published (**Figure 1D**).

**Figure 1.**
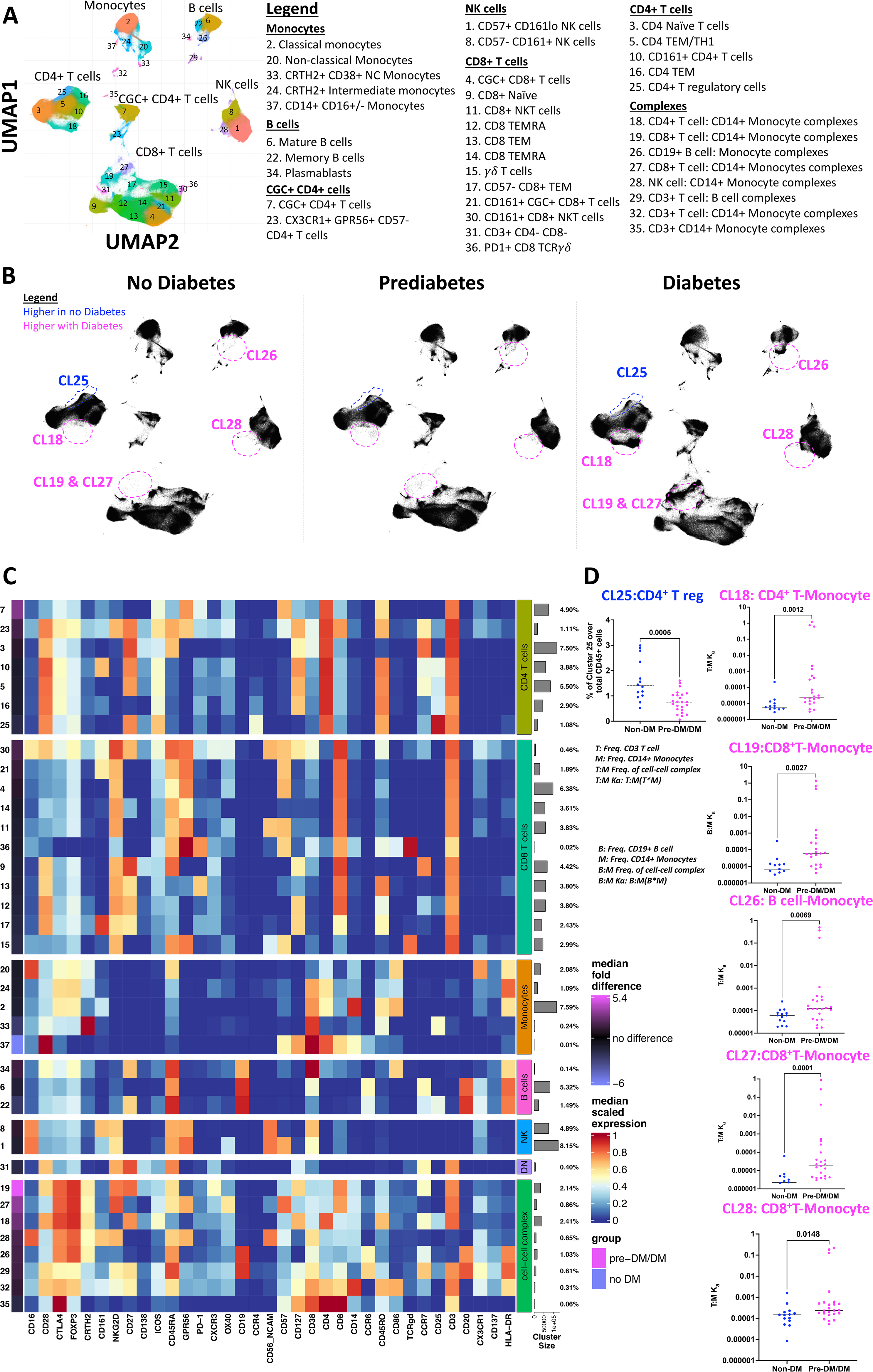
Phenotypic characterization of PBMCs in non-diabetic, prediabetic, and diabetic PLWH highlights differences by metabolic disease categor. (A) UMAP of 1.5 million CD45^+^ cells from the PBMCs of 38 participants with controlled HIV depicting clusters of monocytes, CD4^+^ T cells, CD8^+^ T cells, B cells, and NK cells (B) UMAPs stratified by metabolic disease (no diabetes, prediabetes, and diabetes). For each UMAP, we downsampled to 40000 events per sample from all 38 participants. Clusters 18, 19, 26, 27, and 28 have cell-cell complexes and are significantly higher with prediabetes/diabetes compared to no diabetes. Other clusters that differ by diabetes are included in Table S2 (C) Heat map shows all markers used to define clusters in the UMAPs. The median fold difference legend bar (purple clusters are significantly higher in prediabetics/diabetics and blue are higher in non-diabetic PLWH). Clusters in the heat map are grouped according to the bigger clusters (labels on the right). The percentages indicate the number of cells in that cluster proportional to the total number of cells analyzed. (D) Dot plots show the % CD4^+^ T regulatory cell cluster over total live CD45^+^ and the constant of association between T/B cells and monocytes in the complex clusters 18, 19, 26, 27, and 28 by diabetes status. Statistical analysis by Mann-Whitney test (D). *See Figure S1 and Table S3*.

### T cell-monocyte complexes are increased with glucose intolerance

T-REX workflow, an unbiased machine learning approach, was used to visualize distinct cell populations based on diabetes status and marker enrichment modeling (Barone et al., 2021; Diggins, Greenplate, Leelatian, Wogsland, & Irish, 2017). The UMAP displays clusters that differ between non-diabetic (blue) and prediabetic/diabetic (red) PLWH (**Figure 2A**). All cell-cell complex clusters, except for cluster 3, remained significantly higher in PLWH with diabetes (**Figure 2B**). The expression of CD14^+^ in clusters outside of the monocyte population was confirmed, and other markers like FOXP3 and CTLA4 defined the clusters with cell-cell complexes (**Figure 2C**). Due to their larger proportion, we focused on CD3+T-cell: CD14+monocyte complexes (**Figure 1C**). Additional studies using flow cytometry validated these complexes (**Figure S2A**). Notably, CD3 and CD14 markers were sufficient to help define the T cell-monocyte complexes (**Figure 2Di-iv**).

**Figure 2.**
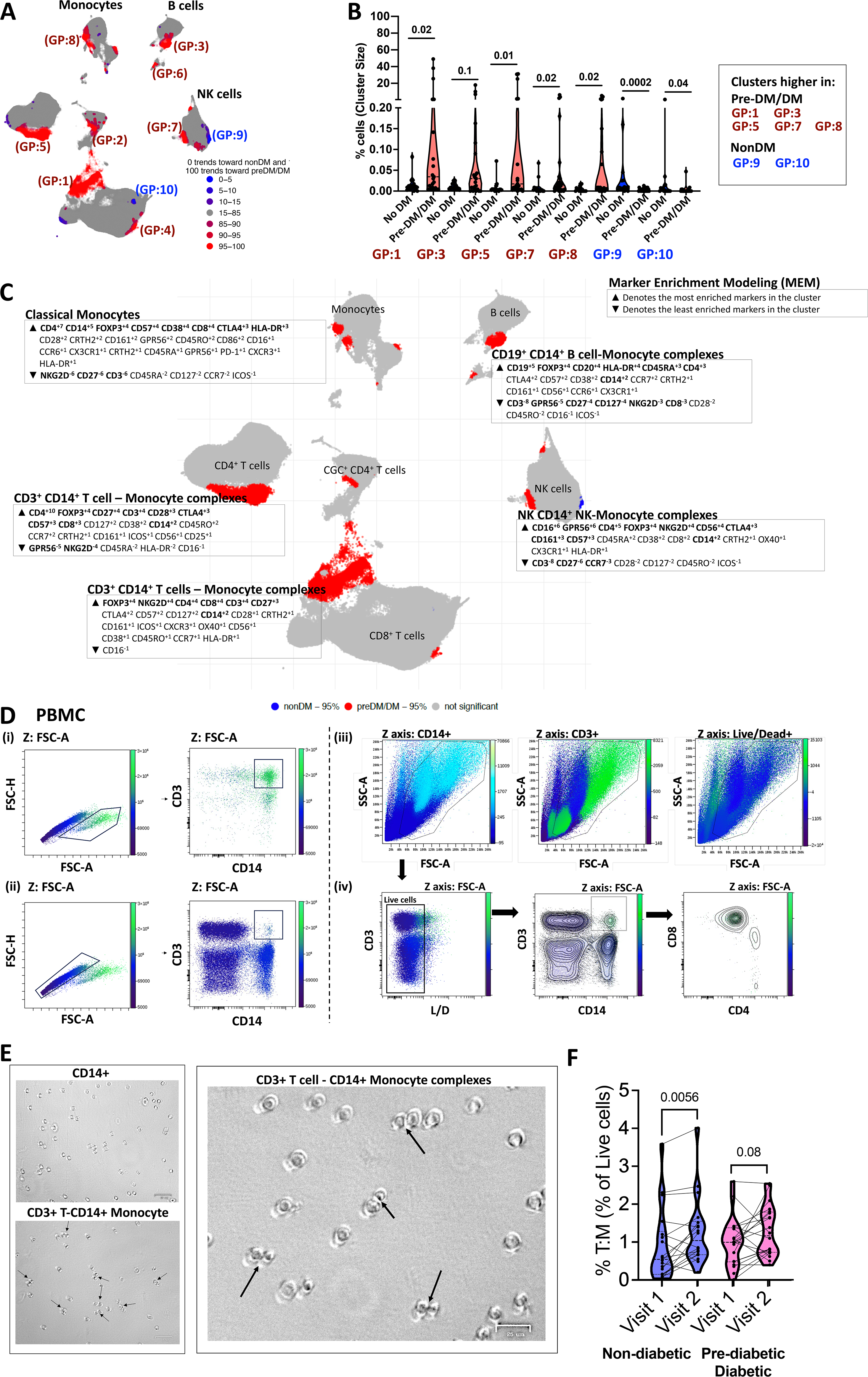
Classical monocytes complexed with T cells, NK cells, and B cells are increased in the peripheral blood of PLWH with glucose intoleranc. (A) Clusters identified by the T-REX algorithm increase and decrease with prediabetes/diabetes (blue is higher in non-diabetics, and red is higher in prediabetic/diabetics) (B) Violin plots show proportions of select subclusters that are significantly different between non-diabetes (blue) and prediabetes/diabetes (maroon) (C) The enrichment scores of markers (increased and decreased) for select clusters are shown over the UMAP adjacent to the clusters. The bold markers^▾^are the most significant among the markers that characterize the clusters. (D) Two-dimensional flow cytometry plot of PBMCs showing FSC-A and FSC-H of cells that comprise complexes (i) and singlets (ii). In section (iii), the FSC-A by SSC-A plots show CD14, CD3, and Live/Dead on the Z-channel. Lastly, (iv) shows live cells (including lymphocytes and monocytes), followed by a two-dimensional plot of CD3^+^ and CD14^+^ cells, and the last plot shows CD4 and CD8 marker expression on the T: M complexes. (E) Bright-field microscopy images of sorted CD14^+^ and CD3^+^ CD14^+^ cells. (F) Violin plot showing % CD3^+^ T cell-CD14^+^ monocyte complexes in longitudinal time points from the same patients 2-3 years apart. To avoid batch effects, all sample comparisons were performed within a single flow cytometry experiment. Statistical analysis by Mann-Whitney test (B) and Wilcoxon matched pair signed ranks test (F). *See Figure S2*. *[Cluster 18: CD4^+^ T cell CD14^+^ Monocyte complex; Cluster 19: CD8^+^ T cell CD14^+^ Monocyte complex; Cluster 25: CD4^+^ T regulatory cell; Cluster 26: B cell-CD14^+^ Monocytes; Cluster 27: CD8^+^ T cell CD14^+^ Monocyte; Cluster 28: NK cell-CD14+ Monocyte; Cluster 29: CD3^+^ T cell B cell; Cluster 32: CD3^+^ T cell CD14^+^ Monocyte]*

We sorted CD14^+^ monocytes, CD3^+^CD4^+^T cells, and CD3^+^T-cell: CD14^+^monocyte complexes and used light microscopy to image the cells. Compared to CD14^+^ monocytes alone, the complexes had a higher proportion of cells with presumed immunological synapses due to the proximity of cells and flattening at the connecting points (**Figure 2E**). In longitudinal analysis, the proportion of CD3^+^ CD14^+^ complexes increased significantly from the first to second visit in non-diabetic PLWH (**Figure 2F**). Finally, we analyzed CD3^+^T-cell: CD14^+^monocyte complexes among the 6 PLWH compared with six HIV-negative individuals with diabetes (**Figure S2A-C**). We observed that metabolic syndrome, regardless of HIV status syndrome, was associated with the formation of these cell-cell complexes (**Figure S2D**).

### T cell-monocyte complexes in PLWH are positively associated with fasting blood glucose and hemoglobin A1C

Based on the observed differences in CD3^+^T-cell: CD14^+^monocyte complexes by diabetes among PLWH, we posited that the cell-cell complexes may be important in glucose intolerance and influenced by factors associated with metabolic disease. To this end, we used partial Spearman rank correlation analysis to assess whether circulating cell-cell complexes identified by mass cytometry (**Figure 1**) were associated with fasting blood glucose and hemoglobin A1C. We adjusted for age, sex, and BMI, two of which were different between PLWH with and without glucose intolerance (**Table S1**). CD8^+^ T cell-CD14^+^ monocyte complexes were associated with fasting blood glucose and hemoglobin. CD4^+^ T cell-CD14^+^ monocyte complexes were positively associated with hemoglobin A1C and triglycerides (**Figure 3A**).

**Figure 3.**
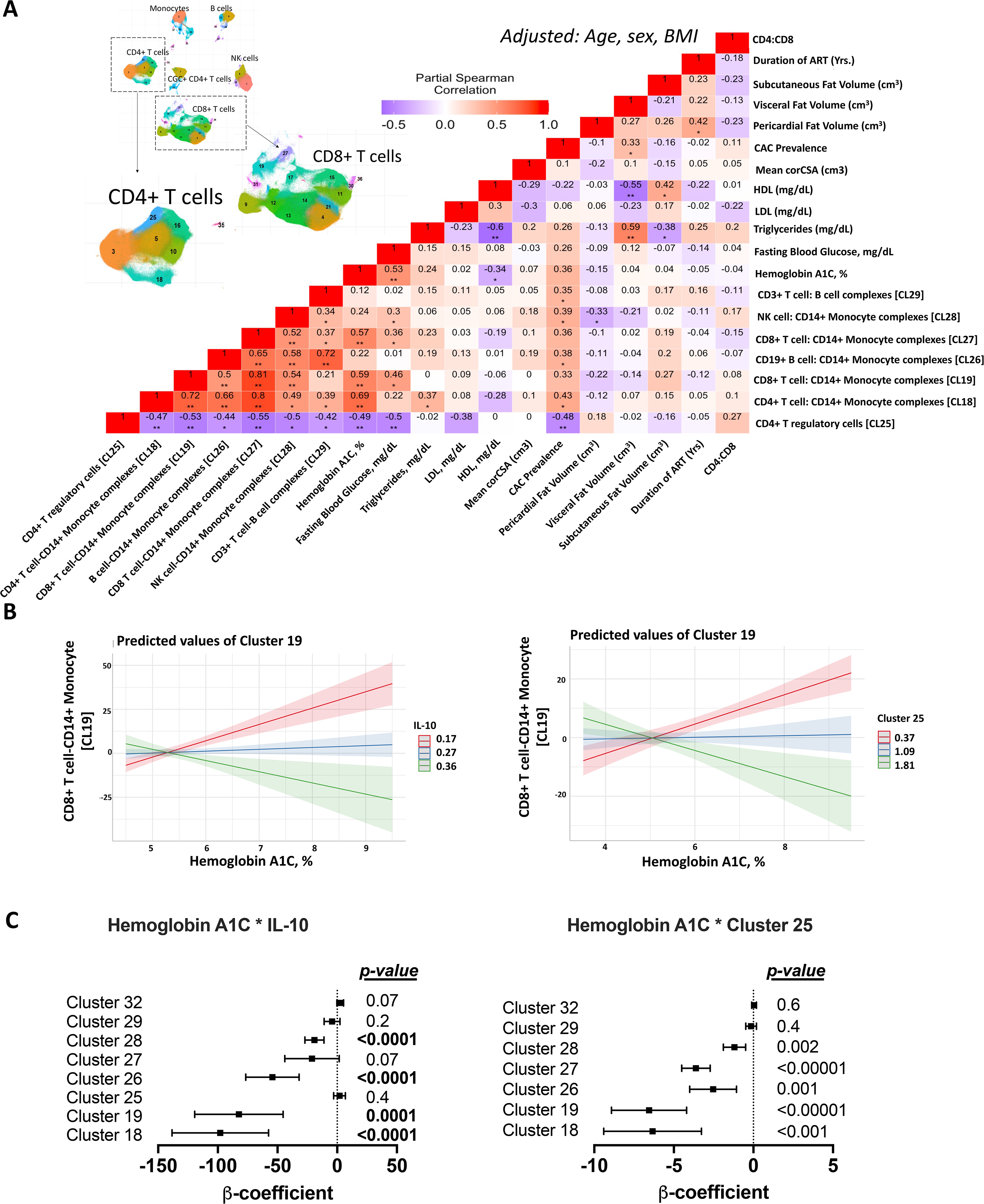
T cell-monocyte complexes are positively associated with blood glucose and negatively with IL-10 and CD4^+^ T regulatory cells in PLWH. (A) Heatmap shows partial Spearman correlation between cell-cell complex clusters, CD4^+^ T regulatory cells from 38 participants as defined by mass cytometry, and hemoglobin A1c, fasting blood glucose adjusted for age, sex, and BMI (* p< 0.05, ** p<0.01). UMAP with clusters from Figure1 is included for reference. (B) Linear regression analysis with cell-cell complexes as the dependent variable and hemoglobin A1C*IL-10 or hemoglobin A1C*Cluster 25 as the independent variables. The line plots depict the relationship between CD8^+^ T cell – CD14^+^ monocyte complexes and hemoglobin A1C, with IL-10 as the interaction term (left) and CD4 T regulatory cells as the interaction term (right). (C) A similar analysis was performed for all cell-cell complex clusters, showing the β coefficients, the 95% confidence intervals, and p-values. *See Figure S3*.

Systemic inflammation is associated with an increased risk of metabolic disease (Hotamisligil, 2006). PLWH on antiretroviral therapy have elevated levels of plasma cytokines at baseline compared to persons without HIV. Among the 38 participants with HIV in this study, there were no differences in select plasma cytokines by metabolic group (**Table S5**). T cell-monocyte complexes were negatively correlated with circulating CD4^+^ T regulatory cells (**Figure 3A**) and plasma IL-10 (**Figure S3**). The negative correlation between T cell-monocyte complexes and circulating CD4^+^ T regulatory cells and IL-10 was modulated by blood glucose levels as determined by hemoglobin A1C as an interaction term (**Figure 3B-C**). This indicates a diminished impact of IL-10 and CD4^+^ T regulatory cells on cell-cell complex formation as glucose increases. Overall, this suggests that there may be a greater tendency for complex formation with metabolic disease, yet the specific roles of the varied immune cells within these complexes remain undetermined.

### T cell-monocyte pairs form stable and dynamic complexes with HIV

Time-lapse imaging (∼5 hours) revealed dynamic T cell-monocyte interactions (**Figure 4A-B, Video 1-2**), with some forming stable complexes (**Figure 4C, D**) and others transient (**Figure 4E**). Control experiments with CD3^+^ singlet T cells and CD14^+^ singlet monocytes showed no complex formation (**Video 3**). We sorted CD3^+^T-cell: CD14^+^monocyte complexes and used TEM to view the interactions. T cells (∼7-12μm with large nuclei) and monocytes (15-18 μm) were identified by morphology (**Figure 4F-i, ii**) (Hossler, 2014; Pavathuparambil Abdul Manaph et al., 2023). We also detected 100nm viral-like particles in these complexes (**Figure 4G**).

**Figure 4.**
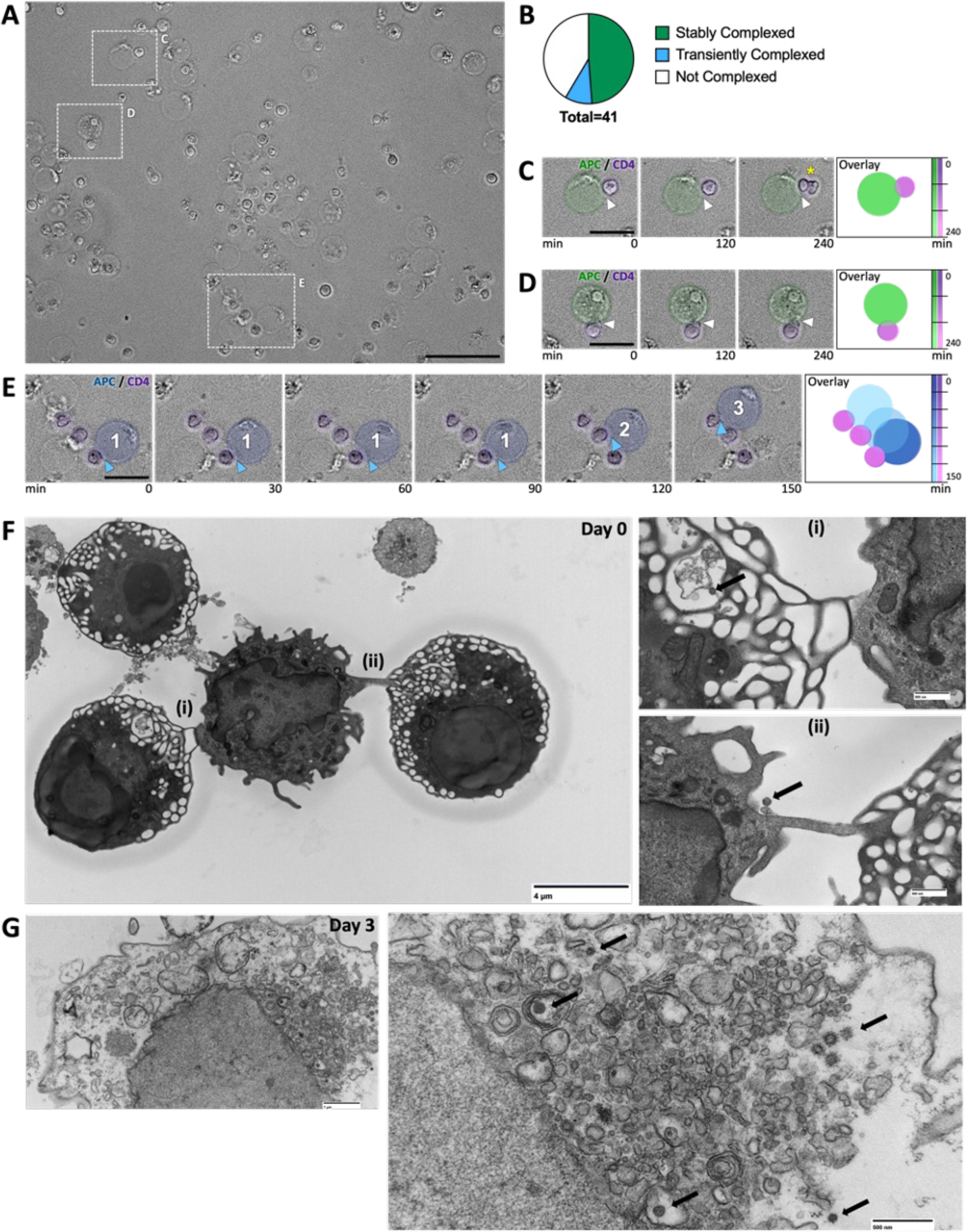
CD3^+^ T cell-CD14^+^ monocyte complexes from PLWH are dynamic. (A) Phase-contrast microscopy of sorted CD3^+^ CD14^+^ T cell-monocyte complexes at time 0. (B) Pie chart shows the percentage of CD14^+^ monocytes that are stably associated with T cells, transiently associated with T cells, or not associated with T cells over 4.5hrs. (C & D) Insets of stable complexes, right-hand panel shows the time overlay and the color code. A yellow asterisk (*) in c marks a T cell that proliferates. Scale bars – purple pseudo color defines T cell and green marks the monocyte. (E) Time series demonstrating transient interactions between CD14^+^ monocyte and three T cells (marked 1,2,3). Blue arrowheads and numbers mark the point of interactions between CD14^+^ monocyte and T cells. (F) TEM of CD3^+^ T cell-CD14^+^ monocyte complexes. Inset highlights ultrastructural cell-cell interactions (i) and (ii) and the presence of 100nm diameter particles (black arrow). (G) TEM of CD3^+^ T cell among sorted CD3^+^ T cell-CD14^+^ monocyte complexes 3 days post-culture. Enlarged image (i) highlighting 100nm diameter particles (black arrow). *Scale bars are 50μm A, 20μm C-E, 4μM F, 500nm F(i), F(ii), G (i), and 1μm G*. See Videos 1-3.

### CD4^+^ T cell-CD14^+^ monocyte complexes are more activated with higher proportions of TH17 cells than singlet CD4^+^ T cells

To better characterize the CD4^+^ T cells complexed with CD14^+^ monocytes, we first analyzed the memory subsets using CCR7 and CD45RO (**Figure 5A**). CD14^+^ monocytes were largely complexed with CD4^+^ TCM and TEM cells (**Figure 5B**). Several markers were used to define activated CD4^+^ T cells (CD137/OX40 and HLADR/CD38) (**Figure 5C**). A significantly higher proportion of activated cells in prediabetic/diabetic PLWH than in non-diabetic CD4^+^ T cells (**Figure 5D, E, left panels**). Focusing on cell-cell complexes, we observed that CD4^+^ T cell-CD14^+^ monocyte complexes had a higher proportion of activated cells (**Figure 5D, right panel**). Irrespective of metabolic status, all cells within cell-cell complexes were HLADR^+^ CD38^+^ (**Figure 5E, right panel**). Circulating activated T cells and cell-cell complexes were correlated with fasting blood glucose (**Figure 5F**). We compared the activation profile between the CD3^+^T-cell: CD14^+^monocyte complexes and singlet T cells and found a higher proportion of CD137^+^OX40^+^ T cells in the cell-cell complexes (**Figure 5G**). Using chemokine receptor markers, we defined CD4^+^ T helper subsets within the CD4^+^ T cells and CD4^+^ T cell-CD14^+^ monocyte complexes (**Figure S1C**). CD3^+^T-cell: CD14^+^monocyte complexes had significantly higher proportions of TH17 cells compared to CD4^+^ T cells (**Figure 5H**). CD4^+^ T cell-CD14^+^ monocyte complexes from prediabetic/diabetic PLWH had a significantly higher proportion of TH2, TH17, and TH1 cells than non-diabetic PLWH (**Figure 5I**). In summary, the T-cell monocyte complexes consist of activated immune cells enriched for TH17 memory subsets.

**Figure 5.**
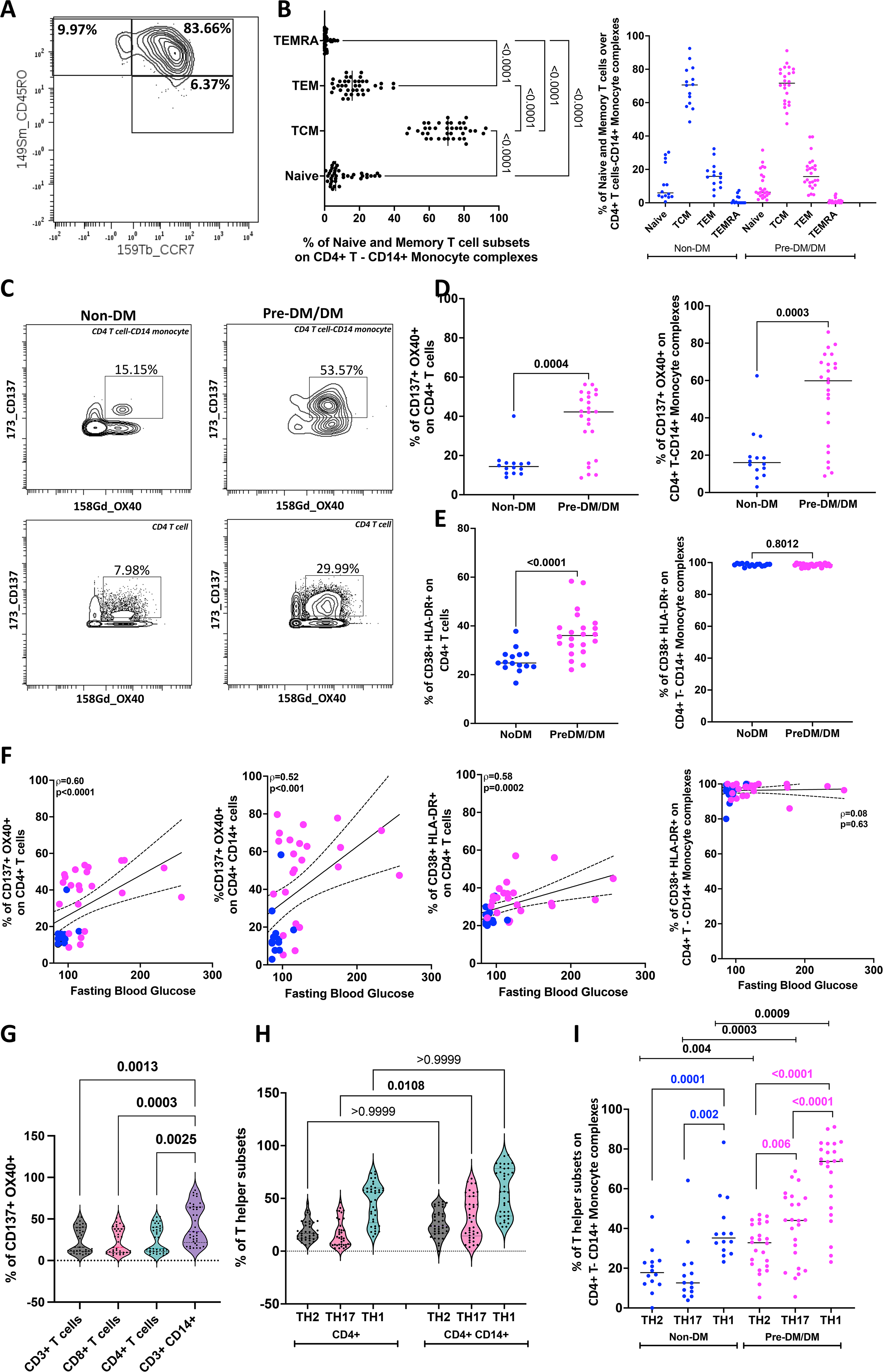
CD4^+^ T cells complexed with CD14^+^ monocytes are more activated with higher proportions of TH17 cells compared to singlet CD4^+^ T cells. (A) Two-dimensional plot of mass cytometry data shows the gating of naïve and memory subsets of CD4^+^ T cells in complex with CD14^+^ monocytes (Naïve, CD45RO^-^ CCR7^+^; TCM, CD45RO^+^ CCR7^+^; TEM CD45RO^+^ CCR7^-^ ; TEMRA CD45RO^-^ CCR7^-^). Gating for CD4^+^ T cells shown in Figure S1A. (B) Dot plots show the proportions of naïve and memory cells in CD4^+^ T cell-CD14^+^ monocyte complexes in all participants (left) and in non-diabetic (n=14) and prediabetic/diabetic PLWH (n=24). (C) Representative plots showing CD137/OX40 on CD4^+^ T cell-CD14^+^ monocyte complexes and CD4^+^ T cells stratified by diabetes. (D) Dot plots show % CD137^+^ OX40^+^ cells on CD4^+^ T cells and on CD4^+^ T cell-CD14^+^ monocyte complexes. (E) Dot plots show % HLA-DR^+^ CD38^+^ cells on CD4^+^ T cells and on CD4^+^ T cell-CD14^+^ monocyte complexes. (F) Correlation plots showing the relationships between fasting blood glucose and % CD137^+^ OX40^+^ cells on CD4^+^ T cells and CD4^+^ T cell-CD14^+^ complexes with. Similar plots of CD38^+^ HLA-DR^+^ expressing cells on CD4^+^ T cells and on CD4^+^ T cell-CD14^+^ monocyte complexes are shown. (G) Violin plots show higher proportions of activated CD137^+^ OX40^+^ cells among CD3^+^ T cell- CD14^+^ monocyte complexes compared to CD3^+^ T cells, CD8^+^ T cells, and CD4^+^ T cells. (H) Violin plots show higher proportions of TH17 cells among CD3^+^ T cell-CD14^+^ monocyte complexes compared to singlet CD4^+^ T cells. (I) PLWH with pre-diabetes/diabetes have a higher proportion of TH2 (CRTH2/CCR4), TH17 (CCR6/CD161), and TH1 (CXCR3) cells as a proportion of CD3^+^ T cell-CD14^+^ monocyte complexes compared to non-diabetic PLWH. Statistical analyses were performed using the Mann-Whitney U test (D-E), Spearman correlation (F), and the Kruskal-Wallis test (G-I).

### T cell-monocyte complexes in PLWH show higher HIV copies and gene expression related to activation and adhesion than singlet T cells and monocytes

We quantified HIV DNA in CD4^+^ T cells, CD14^+^ monocytes, and CD3^+^T-cell: CD14^+^monocyte complexes from six PLWH on ART (selected based on higher proportions of complexes). Representative images show blue droplets (HIV LTR copies) and green droplets (RNAse P copies) (**Figure 6A-C**). A higher count of HIV DNA copies per million cells was observed in CD3^+^T-cell: CD14^+^monocyte complexes compared to paired single CD4^+^ T cells and CD14^+^ monocytes (**Figure 6D-E**).

**Figure 6.**
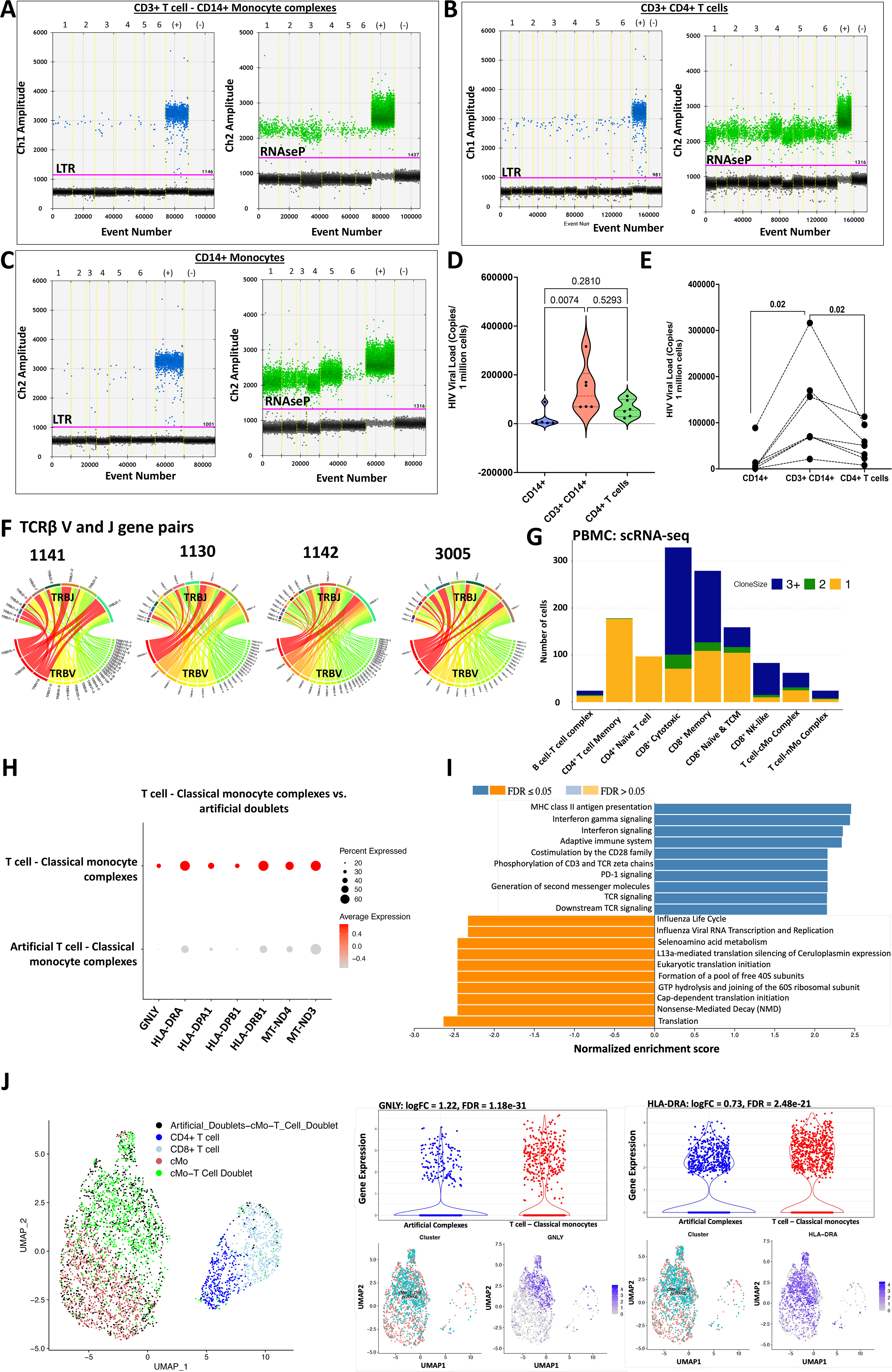
CD3^+^ T cell-CD14^+^ monocyte complexes from PLWH have more copies of HIV compared to singlet CD4^+^ T cells and CD14^+^ monocytes. (A) Representative ddPCR plot showing HIV-LTR (blue droplets) and RNase P (green droplets) copies in sorted CD3^+^ T cell-CD14^+^ monocyte complexes, (B) CD3^+^ CD4^+^ T cells and (C) CD14^+^ monocytes from PLWH. (D) Violin plot shows ddPCR results for HIV quantification from 6 PLWH. (E) The line plot shows HIV viral copies in paired samples. (F) Single CD3^+^ T cell: CD14^+^ monocyte complexes were index-sorted from PBMCs followed by TCR sequencing. The Circos plot shows TCRβ V-J gene pairs of T cells complexed with monocytes from four PLWH (1130, 1141, 1142, and 3005). (G) TCR sequences were obtained from CITE-seq analysis of PBMCs from one individual with many CD3^+^ T cells-CD14^+^ monocyte complexes. The stacked bar chart shows the total number of cells with TCRs and is color-coded based on the clonality of the cells (shared complementarity-determining region 3 (CDR3) sequences with ≥ 2 were considered clonal). (H) Dot plot shows genes that are differentially expressed in T cell-classical monocyte complexes compared to artificial T cell-monocyte complexes from the same scRNA-seq data set. (I) GSEA analysis shows the Reactome pathways enriched by differentially expressed genes that are higher in the T cell-classical monocyte complexes (blue bars) when compared to the artificial complexes (orange bars). (J) UMAP shows artificial complexes and CD3^+^ T cell-CD14^+^ classical monocyte complexes among other T cells (left panel). Violin plots and UMAPs show differential gene expression of GNLY (middle panel) and HLA-DRA (right panel) . Statistical analysis using Kruskal Wallis (D), Wilcoxon test (E). *See Tables S7e*

To determine if the T cells in the complexes are clonally expanded, four PLWH with a larger proportion of cell-cell complexes and paired αβ TCR chains were sequenced from sorted complexes (**Figure 6F**). We used TCRmatch (http://tools.iedb.org/tcrmatch/) to predict the antigen specificity of the clonal TCRs among those identified. The majority of the clonal TCRs (with identical CDR3 >2) were predicted to bind viral antigens, including HIV, in PLWH (**Table S6**). Herpes virus TCRs were predicted in both PLWH and PWoH.

PBMCs from a PLWH with a large proportion of T cell-monocyte complexes as determined by Cytof and flow cytometry were processed for single-cell transcriptomic analysis. In addition to validation based on CD3^+^ and CD14^+^ CITE-seq markers on the same cells, as expected. T cell-monocyte complexes had significantly more reads per cell than all singlet clusters (**Figure S4**). For this sample, classical monocytes complexed with an almost equal representation of T cells with clonal and non-clonal TCRs. Non-classical monocytes, on the other hand, were mostly paired with clonal TCRs (**Figure 6G**). Artificial complexes (a group of singlet monocytes and singlet CD3^+^ T cells combined for analysis) were compared to paired T cell-monocyte complexes from ten participants without HIV who were consented to study immune cells in CVD (**Clinical demographics in Table S7, Figure 6H**). Cell-cell complexes were compared from the same participant for each artificial complex group. Compared to artificial complexes, T cells-monocyte complexes expressed higher levels of GNLY with an overrepresentation of the MHCII antigen presentation and TCR signaling pathways, consistent with an activated inflammatory response (**Figure 6I-J, *Table S8***). The comparisons of the T cells-monocyte complexes with paired artificial complexes in older HIV-negative individuals, showing high inflammation and antigen presentation, suggests that these complexes are not entirely driven by HIV.

### Maintenance of CD3^+^T-cell:CD14^+^monocytes by oxidative phosphorylation

Changes in metabolism can be informative of the functional profile of immune cells (Palmer, Cherry, Sada-Ovalle, Singh, & Crowe, 2016). While immune cells can rely on glucose and mitochondria for energy production, activated cells mostly rely on glycolysis (O’Neill, Kishton, & Rathmell, 2016). We measured the energy dependencies of PBMCs from 15 PLWH (5 non-diabetic and 10 pre-diabetic and diabetic) *ex vivo* using SCENITH (Argüello et al., 2020). Based on the puromycin uptake, CD3^+^T-cell: CD14^+^monocyte complexes and CD14^+^ monocytes were more metabolically active than T cells, given their uptake of puromycin at baseline (**Figure S5A**). The CD3^+^T-cell: CD14^+^monocyte complexes had a higher mitochondrial dependence than the singlet CD14^+^ monocytes (**Figure S5B-D**). We next quantified the proportion of persistent CD3^+^T-cell: CD14^+^monocyte complexes after incubation of PBMCs with metabolic pathway inhibitors (2DG, oligomycin). While the proportion of CD3^+^T-cell: CD14^+^monocyte complexes was unchanged after inhibition of glycolysis with 2DG, there was a decrease with inhibition of oxidative phosphorylation (**Figure S5E**). In summary, CD3^+^T-cell: CD14^+^monocyte complexes use glycolysis and oxidative phosphorylation, though mitochondrial ATP synthesis may play a more significant role in maintaining cell-cell complexes.

## Discussion

Our study reveals that circulating monocytes complexed with T cells are increased in the presence of HIV and glucose intolerance. Many of these complexes remain intact over a 4-hour time-lapse *ex-vivo*, suggesting they can form stable complexes. Furthermore, in one of the Videos, we capture a T cell complex with an antigen-presenting cell dividing (Video 2), suggesting that the complexes are biologically functional. Stable immune complexes with prolonged synapses may contribute to systemic inflammation greater than the contribution of single cells (Friedl & Storim, 2004). Importantly, the cell-cell complexes have increased expression of activation markers and inflammatory gene transcripts. Additionally, there is an inverse association between the cell-cell complexes, plasma IL-10, and CD4^+^ T regulatory cells.

IL-10 is an anti-inflammatory cytokine expressed by several cell types, including macrophages and regulatory T cells.(Moore, de Waal Malefyt, Coffman, & O’Garra, 2001) Several studies, including the Leiden 85-plus study, have shown that immune cells from individuals with metabolic syndrome expressed lower levels of IL-10 upon stimulation.(van Exel et al., 2002) HIV infection is associated with increased expression of IL-10, which in turn can suppress T-cell responses.(Brockman et al., 2009) Over time and with ART, IL-10 expression decreases and may be associated with metabolic disease (Fourman et al., 2020; Werede et al., 2022). In this study, the relationship between the T cell-monocyte complexes and IL-10 and the interaction between hemoglobin A1C and IL-10 reinforces the notion that complexes are increased in the setting of inflammation (de Waal Malefyt, Abrams, Bennett, Figdor, & de Vries, 1991).

In PLWH, replicating and integrated HIV can be detected in circulating monocytes (Lambotte et al., 2000; McElrath, Steinman, & Cohn, 1991; Sonza et al., 2001; Zhu et al., 2002). Although some studies suggest that CD14^lo^ CD16^+^ non-classical monocytes are more prone to HIV infection,(Ellery et al., 2007), we found that HIV is detected in CD4^+^ T cell-CD14^+^ monocyte complexes. Within tissues, macrophages infected with HIV can transmit HIV to CD4^+^ T cells, suggesting these cellular interactions may be an important contributor to T cell loss and the establishment of HIV reservoirs (Carr, Hocking, Li, & Burrell, 1999; Crowe, Zhu, & Muller, 2003; Groot, Welsch, & Sattentau, 2008).

A recent study on immunometabolism of CD4^+^ T cells in the context of HIV showed that the infectivity of the CD4^+^ T cells was more dependent on the metabolic activity of the T cells and less on the activation status (Taylor & Palmer, 2020; Valle-Casuso et al., 2019). They also showed fewer cells with latent HIV infection when they partially inhibited glycolysis using 2DG, suggesting that the steps required for HIV to establish latency are glucose-dependent. In our study, CD3^+^T-cell: CD14^+^monocyte complexes were metabolically active with greater dependence on glucose than oxidative phosphorylation. Forming these cell-cell complexes is an energy-demanding process, and inhibiting oxidative phosphorylation may have been sufficient to affect some of these immunological synapses (Bonifaz, Cervantes-Silva, Ontiveros-Dotor, López-Villegas, & Sánchez-García, 2014).

In summary, we have defined specific features of metabolically active, dynamic T cell-monocyte cell-cell complexes increased with glucose intolerance in the setting of chronic HIV infection. The complex interplay between inflammatory and metabolic disorders makes these cells particularly interesting. Future studies investigating these cells *in vivo* and characterizing HIV within the cell-cell complexes will provide insight into their role in metabolic disease and complications that may arise from this.

## Limitations of the study

This is a cross-sectional study with non-diabetic PWH and pre-diabetic/diabetic PWH (n=14 and n=24, respectively) and differs in variables associated with glucose intolerance, including age, BMI, and waist circumference. While we report an association between the cell-cell complexes and glucose intolerance, we cannot show causality in this study. Therefore, while the inflammatory cell-cell complexes are increased in persons with HIV with increased glucose tolerance and appear to carry HIV, we are currently unable to make conclusions as to whether these cell-cell complexes drive the pathogenesis of the metabolic disease or are a consequence of metabolic disease.

## Author Contributions

Conceptualization, C.N.W., J.R.K., A.K.; Methodology, C.N.W., L.M.O., J.S., J. O., X.Z., Q.S., S.P., J.S., L.M., M.M., S.A.M., R.G., H.B., A. H., E.W., K.N., T.A., Y.R.S., S.A.K., T.T., C.L.G., D.G.H., E.J.P., J.R.K., A.K.; Statistics, C.N.W., A.P., J.S., Q.S., J.S.; Formal Analysis, C.N.W., J.S., Q.S., J.S., J.O., C.M.W., R.G., A.C., S.P. ; Investigation, C.N.W., L.M.O., C.M.W., J.R.K, A.K.; Resources, J.R.K., A.K., S.A.K.; Data Curation, C.N.W., S.B., Q.S., J.S., J. R. K.; Writing – Original Draft, C.N.W.; Writing – Review & Editing original draft., C.N.W., L.M.O., J.S., J. O., X.Z., M.M., S.A.M., H.B., A. H., E.W., S.A.K., T.T., D.G.H., E.J.P., A.K., J.R.K.; Visualization, C.N.W., S.P., J.W., S.B., Q.S., J.S., H.B., A.H.; Supervision, C.N.W, J.R.K., Project Administration, C.N.W., J.R.K.; Funding Acquisition, C.N.W., J.R.K.

## Acknowledgments

Doris Duke CSDA 2021193 (CNW), K23 HL156759 (CNW), Burroughs Wellcome Fund 1021480 (CNW), 1021868.01 (AHJ) and Burroughs Wellcome Fund/ PDEP #1022376 (HKB), R01 DK112262 (JRK), R01HL144941 (AK), R03HL155041 (AK), the Tennessee Center for AIDS Research grant P30 AI110527 (SAM), KL2TR002245 (MM and EMW), K08AR080808 (EMW), The Myositis Association Pilot Award Grant (EMW), the Vanderbilt Flow Cytometry Shared Resource is supported by the Vanderbilt Ingram Cancer Center (P30 CA068485) and the Vanderbilt Digestive Disease Research Center (DK058404), The UNCF/ Bristol-Myers Squibb (UNCF/BMS)-E.E. Just Postgraduate Fellowship in Life sciences Fellowship, NIH Small Research Pilot Subaward to 5R25HL106365-12 from the National Institutes of Health PRIDE Program, DK020593, Vanderbilt Diabetes and Research Training Center for DRTC Alzheimer’s Disease Pilot & Feasibility Program. CZI Science Diversity Leadership grant number 2022-253529 from the Chan Zuckerberg Initiative DAF, an advised fund of the Silicon Valley Community Foundation (AHJ).

The graphical abstract was created using BioRender.

## Declaration of interests

The authors have no competing interests.

## Notes

### Competing Interest Statement

The authors have declared no competing interest.

### Summary of Updates

Revision of text throughout, each figure, and authors.

## References

Abana, C. O., Pilkinton, M. A., Gaudieri, S., Chopra, A., McDonnell, W. J., Wanjalla, C., … Mallal, S. A. (2017). Cytomegalovirus (CMV) Epitope-Specific CD4(+) T Cells Are Inflated in HIV(+) CMV(+) Subjects. J Immunol, 199(9), 3187–3201. doi:10.4049/jimmunol.1700851

Alcaide, M. L., Parmigiani, A., Pallikkuth, S., Roach, M., Freguja, R., Della Negra, M., … Pahwa, S. (2013). Immune activation in HIV-infected aging women on antiretrovirals--implications for age-associated comorbidities: a cross-sectional pilot study. PLoS One, 8(5), e63804. doi:10.1371/journal.pone.0063804

Argüello, R. J., Combes, A. J., Char, R., Gigan, J. P., Baaziz, A. I., Bousiquot, E., … Pierre, P. (2020). SCENITH: A Flow Cytometry-Based Method to Functionally Profile Energy Metabolism with Single-Cell Resolution. Cell Metab, 32(6), 1063–1075.e1067. doi:10.1016/j.cmet.2020.11.007

Arya S, M. D., Kemp SE, Jefferis G. (2019). RANN: Fast Nearest Neighbour Search (Wraps ANN Library) Using L2 Metric. R package version 2.6.1. Retrieved from https://github.com/jefferislab/RANN

Bailin, S. S., Kundu, S., Wellons, M., Freiberg, M. S., Doyle, M. F., Tracy, R. P., … Koethe, J. R. (2022). Circulating CD4+ TEMRA and CD4+ CD28- T cells and incident diabetes among persons with and without HIV. Aids, 36(4), 501–511. doi:10.1097/qad.0000000000003137

Bailin, S. S., McGinnis, K. A., McDonnell, W. J., So-Armah, K., Wellons, M., Tracy, R. P., … Koethe, J. R. (2020). T Lymphocyte Subsets Associated with Prevalent Diabetes in Veterans with and without HIV. J Infect Dis. doi:jiaa069 [pii] 10.1093/infdis/jiaa069 5734988 [pii]

Barone, S. M., Paul, A. G., Muehling, L. M., Lannigan, J. A., Kwok, W. W., Turner, R. B., … Irish, J. M. (2021). Unsupervised machine learning reveals key immune cell subsets in COVID-19, rhinovirus infection, and cancer therapy. Elife, 10. doi:10.7554/eLife.64653

Bolotin, D. A., Poslavsky, S., Mitrophanov, I., Shugay, M., Mamedov, I. Z., Putintseva, E. V., & Chudakov, D. M. (2015). MiXCR: software for comprehensive adaptive immunity profiling. Nat Methods, 12(5), 380–381. doi:10.1038/nmeth.3364

Bonifaz, L., Cervantes-Silva, M., Ontiveros-Dotor, E., López-Villegas, E., & Sánchez-García, F. (2014). A Role For Mitochondria In Antigen Processing And Presentation. Immunology, 144(3), 461–471. doi:10.1111/imm.12392

Brockman, M. A., Kwon, D. S., Tighe, D. P., Pavlik, D. F., Rosato, P. C., Sela, J., … Kaufmann, D. E. (2009). IL-10 is up-regulated in multiple cell types during viremic HIV infection and reversibly inhibits virus-specific T cells. Blood, 114(2), 346–356. doi:10.1182/blood-2008-12-191296

Burel, J. G., Pomaznoy, M., Lindestam Arlehamn, C. S., Weiskopf, D., da Silva Antunes, R., Jung, Y., … Peters, B. (2019). Circulating T cell-monocyte complexes are markers of immune perturbations. Elife, 8. doi:10.7554/eLife.46045

Carr, J. M., Hocking, H., Li, P., & Burrell, C. J. (1999). Rapid and efficient cell-to-cell transmission of human immunodeficiency virus infection from monocyte-derived macrophages to peripheral blood lymphocytes. Virology, 265(2), 319–329. doi:10.1006/viro.1999.0047

Crowe, S., Zhu, T., & Muller, W. A. (2003). The contribution of monocyte infection and trafficking to viral persistence, and maintenance of the viral reservoir in HIV infection. J Leukoc Biol, 74(5), 635–641. doi:10.1189/jlb.0503204

de Waal Malefyt, R., Abrams, J., Bennett, B., Figdor, C. G., & de Vries, J. E. (1991). Interleukin 10(IL-10) inhibits cytokine synthesis by human monocytes: an autoregulatory role of IL-10 produced by monocytes. J Exp Med, 174(5), 1209–1220. doi:10.1084/jem.174.5.1209

Diggins, K. E., Greenplate, A. R., Leelatian, N., Wogsland, C. E., & Irish, J. M. (2017). Characterizing cell subsets using marker enrichment modeling. Nat Methods, 14(3), 275–278. doi:10.1038/nmeth.4149

Ellery, P. J., Tippett, E., Chiu, Y. L., Paukovics, G., Cameron, P. U., Solomon, A., … Crowe, S. M. (2007). The CD16+ monocyte subset is more permissive to infection and preferentially harbors HIV-1 in vivo. J Immunol, 178(10), 6581–6589.

Ellis B, H. P., Hahne F, Le Meur N, Gopalakrishnan N, Spidlen J, Jiang M, Finak G. (2022). FlowCore: flowCore: Basic structures for flow cytometry data_. R package version 2.8.0. Retrieved from https://github.com/RGLab/flowCore

Fourman, L. T., Saylor, C. F., Cheru, L., Fitch, K., Looby, S., Keller, K., … Lo, J. (2020). Anti-Inflammatory Interleukin 10 Inversely Relates to Coronary Atherosclerosis in Persons With Human Immunodeficiency Virus. J Infect Dis, 221(4), 510–515. doi:10.1093/infdis/jiz254

Friedl, P., & Storim, J. (2004). Diversity in immune-cell interactions: states and functions of the immunological synapse. Trends Cell Biol, 14(10), 557–567. doi:10.1016/j.tcb.2004.09.005

Gherardini, F. (2022). Premessa: R package for pre-processing of flow and mass cytometry data. Retrieved from https://github.com/ParkerICI/premessa

Gil-Manso, S., Miguens Blanco, I., López-Esteban, R., Carbonell, D., López-Fernández, L. A., West, L., … Pion, M. (2021). Comprehensive Flow Cytometry Profiling of the Immune System in COVID-19 Convalescent Individuals. Front Immunol, 12, 793142. doi:10.3389/fimmu.2021.793142

Grome, H. N., Barnett, L., Hagar, C. C., Harrison, D. G., Kalams, S. A., & Koethe, J. R. (2017). Association of T Cell and Macrophage Activation with Arterial Vascular Health in HIV. AIDS Res Hum Retroviruses, 33(2), 181–186. doi:10.1089/AID.2016.0113

Groot, F., Welsch, S., & Sattentau, Q. J. (2008). Efficient HIV-1 transmission from macrophages to T cells across transient virological synapses. Blood, 111(9), 4660–4663. doi:10.1182/blood-2007-12-130070

Grün, D., Kester, L., & van Oudenaarden, A. (2014). Validation of noise models for single-cell transcriptomics. Nat Methods, 11(6), 637–640. doi:10.1038/nmeth.2930

Hao, Y., Hao, S., Andersen-Nissen, E., Mauck, W. M., Zheng, S., Butler, A., … Satija, R. (2021). Integrated analysis of multimodal single-cell data. Cell, 184(13), 3573–3587.e3529. doi:10.1016/j.cell.2021.04.048

Hertzman, R. J., Deshpande, P., Leary, S., Li, Y., Ram, R., Chopra, A., … Phillips, E. J. (2021). Visual Genomics Analysis Studio as a Tool to Analyze Multiomic Data. Front Genet, 12, 642012. doi:10.3389/fgene.2021.642012

Hossler, F. (2014). Ultrastructure Atlas of Human Tissues: John Wiley & Sons.

Hotamisligil, G. S. (2006). Inflammation and metabolic disorders. Nature, 444(7121), 860–867. doi:10.1038/nature05485

Hsue, P. Y., & Waters, D. D. (2019). HIV infection and coronary heart disease: mechanisms and management. Nat Rev Cardiol, 16(12), 745–759. doi:10.1038/s41569-019-0219-9

Islam, S., Zeisel, A., Joost, S., La Manno, G., Zajac, P., Kasper, M., … Linnarsson, S. (2014). Quantitative single-cell RNA-seq with unique molecular identifiers. Nat Methods, 11(2), 163–166. doi:10.1038/nmeth.2772

J, M. (2022). Uwot: The Uniform Manifold Approximation and Projection (UMAP) Method for Dimensionality Reduction_. R package version 0.1.14. Retrieved from https://github.com/jlmelville/uwot

Kelly, S. T. (2020). Leiden: R implementation of the Leiden algorithm. R package version 0.4.3 2022 R package version 0.4.3. Retrieved from https://github.com/TomKellyGenetics/leiden

Kivioja, T., Vähärautio, A., Karlsson, K., Bonke, M., Enge, M., Linnarsson, S., & Taipale, J. (2011). Counting absolute numbers of molecules using unique molecular identifiers. Nat Methods, 9(1), 72–74. doi:10.1038/nmeth.1778

Kundu, S., Freiberg, M. S., Tracy, R. P., So-Armah, K. A., Koethe, J. R., Duncan, M. S., … Study, V. A. C. (2022). Circulating T Cells and Cardiovascular Risk in People With and Without HIV Infection. J Am Coll Cardiol, 80(17), 1633–1644. doi:10.1016/j.jacc.2022.08.756

Lam, J., Katti, P., Biete, M., Mungai, M., AshShareef, S., Neikirk, K., … Hinton, A. (2021). A Universal Approach to Analyzing Transmission Electron Microscopy with ImageJ. Cells, 10(9). doi:10.3390/cells10092177

Lambotte, O., Taoufik, Y., de Goër, M. G., Wallon, C., Goujard, C., & Delfraissy, J. F. (2000). Detection of infectious HIV in circulating monocytes from patients on prolonged highly active antiretroviral therapy. J Acquir Immune Defic Syndr, 23(2), 114–119. doi:10.1097/00126334-200002010-00002

McElrath, M. J., Steinman, R. M., & Cohn, Z. A. (1991). Latent HIV-1 infection in enriched populations of blood monocytes and T cells from seropositive patients. J Clin Invest, 87(1), 27–30. doi:10.1172/JCI114981

Moore, K. W., de Waal Malefyt, R., Coffman, R. L., & O’Garra, A. (2001). Interleukin-10 and the interleukin-10 receptor. Annu Rev Immunol, 19, 683–765. doi:10.1146/annurev.immunol.19.1.683

Neikirk, K., Vue, Z., Katti, P., Rodriguez, B. I., Omer, S., Shao, J., … Hinton, A. O. (2023). Systematic Transmission Electron Microscopy-Based Identification and 3D Reconstruction of Cellular Degradation Machinery. Adv Biol (Weinh), e2200221. doi:10.1002/adbi.202200221

O’Neill, L. A., Kishton, R. J., & Rathmell, J. (2016). A guide to immunometabolism for immunologists. Nat Rev Immunol, 16(9), 553–565. doi:10.1038/nri.2016.70

Palmer, C. S., Cherry, C. L., Sada-Ovalle, I., Singh, A., & Crowe, S. M. (2016). Glucose Metabolism in T Cells and Monocytes: New Perspectives in HIV Pathogenesis. EBioMedicine, 6, 31–41. doi:10.1016/j.ebiom.2016.02.012

Pavathuparambil Abdul Manaph, N., Ltaief, S. M., Nour-Eldine, W., Ponraj, J., Agcaoili, J., Mansour, S., & Al-Shammari, A. R. (2023). An optimized protocol for the preparation of blood immune cells for transmission electron microscopy. Micron, 173, 103517. doi:10.1016/j.micron.2023.103517

Sivakumar, S., Abu-Shah, E., Ahern, D. J., Arbe-Barnes, E. H., Jainarayanan, A. K., Mangal, N., … Dustin, M. L. (2021). Activated Regulatory T-Cells, Dysfunctional and Senescent T-Cells Hinder the Immunity in Pancreatic Cancer. Cancers (Basel), 13(8). doi:10.3390/cancers13081776

Sonza, S., Mutimer, H. P., Oelrichs, R., Jardine, D., Harvey, K., Dunne, A., … Crowe, S. M. (2001). Monocytes harbour replication-competent, non-latent HIV-1 in patients on highly active antiretroviral therapy. AIDS, 15(1), 17–22. doi:10.1097/00002030-200101050-00005

Taylor, H. E., & Palmer, C. S. (2020). CD4 T Cell Metabolism Is a Major Contributor of HIV Infectivity and Reservoir Persistence. Immunometabolism, 2(1). doi:10.20900/immunometab20200005

Temu, T. M., Polyak, S. J., Zifodya, J. S., Wanjalla, C. N., Koethe, J. R., Masyuko, S., … Farquhar, C. (2020). Endothelial Dysfunction Is Related to Monocyte Activation in Antiretroviral-Treated People With HIV and HIV-Negative Adults in Kenya. Open Forum Infect Dis, 7(10), ofaa425. doi:10.1093/ofid/ofaa425

Temu, T. M., Wagoner, J., Masyuko, S., O’Connor, A., Zifodya, J. S., Macharia, P., … Polyak, S. J. (2021). Central obesity is a contributor to systemic inflammation and monocyte activation in virally suppressed adults with chronic HIV in Kenya. AIDS, 35(11), 1723–1731. doi:10.1097/QAD.0000000000002956

Valle-Casuso, J. C., Angin, M., Volant, S., Passaes, C., Monceaux, V., Mikhailova, A., … Sáez-Cirión, A. (2019). Cellular Metabolism Is a Major Determinant of HIV-1 Reservoir Seeding in CD4. Cell Metab, 29(3), 611–626.e615. doi:10.1016/j.cmet.2018.11.015

van Exel, E., Gussekloo, J., de Craen, A. J., Frölich, M., Bootsma-Van Der Wiel, A., Westendorp, R. G., & Study, L. P. (2002). Low production capacity of interleukin-10 associates with the metabolic syndrome and type 2 diabetes : the Leiden 85-Plus Study. Diabetes, 51(4), 1088–1092. doi:10.2337/diabetes.51.4.1088

Wang, J., Duncan, D., Shi, Z., & Zhang, B. (2013). WEB-based GEne SeT AnaLysis Toolkit (WebGestalt): update 2013. Nucleic Acids Res, 41(Web Server issue), W77–83. doi:10.1093/nar/gkt439

Wanjalla, C. N., Mashayekhi, M., Bailin, S., Gabriel, C. L., Meenderink, L. M., Temu, T., … Koethe, J. R. (2021). Anticytomegalovirus CD4 T Cells Are Associated With Subclinical Atherosclerosis in Persons With HIV. Arterioscler Thromb Vasc Biol, Apr;41(4):1459–1473., ATVBAHA120315786. doi:10.1161/ATVBAHA.120.315786

Wanjalla, C. N., McDonnell, W. J., Barnett, L., Simmons, J. D., Furch, B. D., Lima, M. C., … Koethe, J. R. (2019). Adipose Tissue in Persons With HIV Is Enriched for CD4. Front Immunol, 10, 408. doi:10.3389/fimmu.2019.00408

Wanjalla, C. N., McDonnell, W. J., Ram, R., Chopra, A., Gangula, R., Leary, S., … Koethe, J. R. (2021a). Single-cell analysis shows that adipose tissue of persons with both HIV and diabetes is enriched for clonal, cytotoxic, and CMV-specific CD4+ T cells. Cell Rep Med, 2(2), 100205. doi:10.1016/j.xcrm.2021.100205

Wanjalla, C. N., McDonnell, W. J., Ram, R., Chopra, A., Gangula, R., Leary, S., … Koethe, J. R. (2021b). Single-cell analysis shows that adipose tissue of persons with both HIV and diabetes is enriched for clonal, cytotoxic, and CMV-specific CD4+ T cells. Cell Rep Med, 2(2), 100205. doi:10.1016/j.xcrm.2021.100205

Werede, A. T., Terry, J. G., Nair, S., Temu, T. M., Shepherd, B. E., Bailin, S. S., … Wanjalla, C. N. (2022). Mean Coronary Cross-Sectional Area as a Measure of Arterial Remodeling Using Noncontrast CT Imaging in Persons With HIV. J Am Heart Assoc, 11(23), e025768. doi:10.1161/JAHA.122.025768

Zhu, T., Muthui, D., Holte, S., Nickle, D., Feng, F., Brodie, S., … Corey, L. (2002). Evidence for human immunodeficiency virus type 1 replication in vivo in CD14(+) monocytes and its potential role as a source of virus in patients on highly active antiretroviral therapy. J Virol, 76(2), 707–716. doi:10.1128/jvi.76.2.707-716.2002

