## Supplemental Figures for "CD3^+^ T-cell: CD14^+^monocyte complexes are dynamic and increased with HIV and glucose intolerance"

Running Title (54 characters): Cell complexes in HIV, diabetes, and Hypertension

Keywords (5): HIV, CD3^+^ T cell-CD14^+^ monocyte complexes, doublets, diabetes, reservoir

**Table S1. Clinical Demographics of HIV cohort**

|  | **N** | | **Non-Diabetic**  **PWH N=14** | | **Pre-diabetic/Diabetic PWH N=24** | **P-value** |
| --- | --- | --- | --- | --- | --- | --- |
| Age, Yrs. | 38 | 39 [36, 53] | | 49 [40, 56] | | 0.1 |
| Race | 38 |  | |  | | 0.3 |
| African American |  | 0.36 ^5^⁄_14_ | | 0.58 ^14^⁄_24_ | |  |
| Caucasian |  | 0.57 ^8^⁄_14_ | | 0.42 ^10^⁄_24_ | |  |
| Other |  | 0.07 ^1^⁄_14_ | | 0.00 ^0^⁄_24_ | |  |
| Sex : male | 38 | 0.64 ^9^⁄_14_ | | 0.62 ^15^⁄_24_ | | 0.9 |
| Body mass index (kg/m^2^) | 38 | 31.4 [29.8, 34.3] | | 34.4 [30.7, 39.0] | | **<0.05** |
| Hip circumference (cm) | 37 | 107 [104, 116] | | 113 [108, 125] | | **<0.05** |
| Waist circumference (cm) | 37 | 102 [96, 108] | | 109 [104, 117] | | **0.01** |
| Smoker status : | 38 | 0.00 ^0^⁄_14_ | | 0.04 ^1^⁄_24_ | | 0.2 |
| NO |  | 0.64 ^9^⁄_14_ | | 0.83 ^20^⁄_24_ | |  |
| YES |  | 0.36 ^5^⁄_14_ | | 0.12 ^3^⁄_24_ | |  |
| **Laboratory Measures** | | | | | | |
| Fasting Blood glucose, mg/dL | 37 | 89 [86, 95] | | 116 [101, 127] | | **<0.001** |
| LDL, mg/dL | 37 | 102 [84, 118] | | 93 [77, 122] | | 0.8 |
| Cholesterol, mg/dL | 38 | 170 [161, 196] | | 180 [146, 209] | | 0.7 |
| HDL, mg/dL | 38 | 46 [38, 52] | | 41 [34, 51] | | 0.6 |
| Triglycerides, mg/dL | 38 | 87 [74, 232] | | 160 [95, 237] | | 0.2 |
| HsCRP | 37 | 2.5 [1.7, 4.2] | | 3.0 [1.5, 4.8] | | 0.5 |
| **HIV-Related Labs** | | | | | | |
| CD4 at ART start (cells/ml) | 37 | 453 [354, 592] | | 470 [392, 606] | | 0.8 |
| CD4 T cell at enrollment (cells/ml) | 38 | 869 [683, 990] | | 935 [717, 1184] | | 0.4 |
| Cell associated DNA, cells per million copies | 19 | 1385 [669, 1601] | | 347 [178, 1195] | | 0.1 |
| Cell associated RNA, cells per million copies | 19 | 1423 [1260, 3320] | | 656 [171, 2058] | | 0.1 |
| Current ART : | 38 |  | |  | | 0.7 |
| Other |  | 0.07 ^1^⁄_14_ | | 0.12 ^3^⁄_24_ | |  |
| Protease inhibitors & NRTIs |  | 0.14 ^2^⁄_14_ | | 0.04 ^1^⁄_24_ | |  |
| Integrase inhibitors & NRTIs |  | 0.57 ^8^⁄_14_ | | 0.54 ^13^⁄_24_ | |  |
| Protease inhibitors & NRTIs |  | 0.07 ^1^⁄_14_ | | 0.17 ^4^⁄_24_ | |  |
| NNRTI and NRTIs |  | 0.14 ^2^⁄_14_ | | 0.12 ^3^⁄_24_ | |  |
| Duration ART (Yrs) | 37 | 8.3 [4.3, 17.5] | | 9.0 [4.4, 14.4] | | 1 |
| Hepatitis C ab status | 38 | 0.07 ^1^⁄_14_ | | 0.08 ^2^⁄_24_ | | 0.9 |
| **CT Imaging** | | | | | | |
| Pericardial Fat Volume (cm3) | 38 | 78 [58, 127] | | 64 [44, 164] | | 0.6 |
| Visceral Fat Volume (cm3) | 38 | 147 [120, 180] | | 191 [129, 226] | | 0.1 |
| Liver mean hu | 38 | 61.3 [57.0, 65.1] | | 61.0 [56.0, 64.4] | | 1 |
| Subcutaneous Fat Volume (cm3) | 38 | 395 [351, 426] | | 444 [287, 583] | | 0.7 |
| CAC Prevalence : Yes | 38 | 0.00 ^0^⁄_14_ | | 0.33 ^8^⁄_24_ | | **0.02** |
| The statistics represent the median, lower and upper quartile for continuous variables.   N is the number of non-missing values.  Tests used: Wilcoxon test for continuous variables; Pearson chi-squared test for categorical variables | | | | | | |

**Table S2. Reagents used in experimental methods**

| **CYTOF Reagents** | | | |
| --- | --- | --- | --- |
| **Target** | **Metal Tag** | **Tag Isotope** | **Source or reference** |
| CD45 | Y | 89 | Fluidigm |
| CCR6 | Praseodymium (Pr) | 141 | Fluidigm |
| CD57 | Neodymium (Nd) | 142 | Fluidigm |
| CD127 | Neodymium (Nd) | 143 | Fluidigm |
| CD38 | Neodymium (Nd) | 144 | Fluidigm |
| CD4 | Neodymium (Nd) | 145 | Fluidigm |
| CD8a | Neodymium (Nd) | 146 | Fluidigm |
| CD14 M5E2 | Neodymium (Nd) | 148 | Fluidigm |
| CD45RO | Samarium (Sm) | 149 | Fluidigm |
| CD86 | Neodymium | 150 | Fluidigm |
| ICOS | Europium (Eu) | 151 | Fluidigm |
| TCRγδ | Samarium (Sm) | 152 | Fluidigm |
| CD45RA | Europium (Eu) | 153 | Fluidigm |
| GPR56 | Samarium (Sm) | 154 | CIC core |
| PD-1 | Gadolinium (Gd) | 155 | Fluidigm |
| CXCR3 | Gadolinium (Gd) | 156 | Fluidigm |
| OX40 | Gadolinium (Gd) | 158 | Fluidigm |
| CCR7 | Terbium (Tb) | 159 | Fluidigm |
| CD28 | Gadolinium (Gd) | 160 | Fluidigm |
| CD152/CTLA4 | Dysprosium (Dy) | 161 | Fluidigm |
| FoxP3 | Dysprosium (Dy) | 162 | Fluidigm |
| CRTH2 | Dysprosium (Dy) | 163 | Fluidigm |
| CD161 | Dysprosium (Dy) | 164 | Fluidigm |
| CD19 | Holmium(Ho) | 165 | Fluidigm |
| NKG2D | Erbium (Er) | 166 | Fluidigm |
| CD27 | Erbium (Er) | 167 | Fluidigm |
| CD138 | Erbium (Er) | 168 | Fluidigm |
| CD25 | Thulium (TM) | 169 | Fluidigm |
| CD3 | Erbium (Er) | 170 | Fluidigm |
| CD20 | Ytterbium (Yb) | 171 | Fluidigm |
| CX3CR1 | Ytterbium (Yb) | 172 | CIC core |
| CD137 | Ytterbium (Yb) | 173 | Fluidigm |
| HLA-Dr | Lutetium (Lu) | 174 | Fluidigm |
| CCR4 | Lutetium (Lu) | 175 | Fluidigm |
| CD56/NCAM | Ytterbium (Yb) | 176 | Fluidigm |
| Nuc acid --Ir | Iridium (Ir) | 191/193 | Fluidigm |
| Cisplatin | Platinum (Pt) | 195 | Fluidigm |
| CD16 | Bismuth (Bi) | 209 | Fluidigm |
| **Flow Cytometry Reagents**  **BD ARIA** | | | |
| **Target** | **Fluorescent Tag** | **Source or Reference** | **Catalog number** |
| CD3 (Clone SK7) | BV786 | BD Biosciences | #563800 |
| CD4 (Clone RPA-T4) | PcPCy5.5 | BD Biosciences | #560650 |
| CD8 (Clone PRA-T8) | A700 | BD Biosciences | #557945 |
| CCR7 (Clone 150503) | BV421 | BD Biosciences | #562555 |
| CD45RO (Clone UCHL1) | PECF594 | BD Biosciences | #562299 |
| LIVE/DEAD Fixable Aqua | N/A | ThermoFisher | #L34957 |
| CD57 (Lot 4182924) | FITC | BD Pharmingen | # 555619 |
| CX3CR1(Clone 2A9-1) | PE | BD Biosciences | #565796 |
| GPR56 (Clone CG4) | PECY7 | BioLegend | #358205 |
| CD14 (Clone M5E2) | APC | BD Biosciences | #561383 |
| CD19 (Clone HIB19) | V500 | BD Biosciences | #561121 |
| **CYTEK** | | | |
| GPR56 (Clone 4C3) | APC | BioLegend | #391905 |
| CCR7 (Clone G043H7) | Spark NIR 685 | BioLegend | #353257 |
| CD38 (Clone HIT2) | AF700 | BD Biosciences | #560676 |
| KLRG1 (Clone 2F1-KLRG1) | APC CY7 | BioLegend | #138425 |
| CD14 (Clone 63D3) | APC Fire 810 | BioLegend | #367155 |
| CX3CR1 (Clone 2A9-1) | PE | BD Biosciences | #565796 |
| CD45RO (Clone UCHL1) | PE-CF594 | BD Biosciences | #562299 |
| CXCR3 (Clone G025H7) | PE-CY5 | BioLegend | #353755 |
| PD1 (Clone EH12.2H7) | PE Fire 700 | BioLegend | #329918 |
| CD27 (Clone O323) | PE-CY7 | BioLegend | #302859 |
| Puromycin AF488 | PE Fire 810 | Millipore, | #MABE343-AF488 |
| CD16 PerCP | PerCP | BioLegend | #302030 |
| L/D Zombie Violet | BB660 | BioLegend | #423113 |
| CD3 BV480 (Clone UCHT1) | BV480 | BD Biosciences | #566166 |
| CD19 BV510 (Clone HIB19) | BB510 | BioLegend | #302241 |
| CD8 BV570 (Clone RPA-T8) | BB570 | BioLegend | #301037 |
| CD4 BV650 (Clone RPA-T4) | BV650 | BioLegend | #300535 |
| CD28 BV711 (Clone CD28.2) | BV711 | BioLegend | #302947 |
| CXCR5 BV785 (Clone J252D24) | BV785 | BioLegend | #356935 |
| **CITE-seq Reagents** |  |  |  |
| **Target** | **Barcode** | **Source or Reference** | **Catalog number** |
| TotalSeq™-anti-human CD3 (Clone UCHT1) | CTCATTGTAACTCCT | BioLegend | #C0034 |
| TotalSeq™-anti-human CD4 (Clone RPA-T4) | TGTTCCCGCTCAACT | BioLegend | #C0072 |
| TotalSeq™-anti-human CD14 Antibody (Clone M5E2) | TCTCAGACCTCCGTA | BioLegend | #C0081 |
| TotalSeq™-anti-human CD16 Antibody (Clone 3G8) | AAGTTCACTCTTTGC | BioLegend | #C0083 |
| TotalSeq™-anti-human CD11b Antibody (Clone ICRF44) | GACAAGTGATCTGCA | BioLegend | #C0161 |
| TotalSeq™-anti-human CD11c Antibody (Clone S-HCL-3) | TACGCCTATAACTTG | BioLegend | #C0053 |
| TotalSeq™-anti-human CD69 Antibody (Clone FN50) | GTCTCTTGGCTTAAA | BioLegend | #C0146 |
| TotalSeq™-anti-human CD19 Antibody (Clone HIB19) | CTGGGCAATTACTCG | BioLegend | #C0050 |
| TotalSeq™-anti-human CD20 Antibody (Clone 2H7) | TTCTGGGTCCCTAGA | BioLegend | #C0100 |
| TotalSeq™- anti-human CD21 Antibody (clone Bu32) | AACCTAGTAGTTCGG | BioLegend | #C0181 |
| **Energy Metabolism Profiling with SCENITH** | | | |
| Anti-puromycin (Clone 12D10) | AF488 | Millipore Sigma | #MABE343 |
| Puromycin solution | N/A | Sigma Aldrich | #P8833-10MG |
| Oligomycin A | N/A | Sigma Aldrich | #75351-5MG |
| 2-deoxy-D-glucose 2mM | N/A | Sigma Aldrich | #D8375 |
| **Software and Algorithms** | | | |
| R 4.1.2 | The R project for Statistical Computing | https://www.r-project.org/ |  |
| Cytobank Vanderbilt | Cytobank | Vanderbilt.cytobank.org |  |
| GraphPad Prism version 9.5.0 | GraphPad Software | https://www.graphpad.com |  |
| FlowJo V | Tree Star | https://www.flowjo.com |  |
| TCRmatch | IEDB | http://tools.iedb.org/tcrmatch/ |  |
| VGAS | IIID | https://www.iiid.com.au/software/vgas |  |

**Table S3. Classification of cell:cell complexes by different modalities**

| **Modality** | **Definition of cell-cell complexes** |
| --- | --- |
| Flow cytometry | Identification of cell:cell complexes using CD3 and CD14 antibodies, which ordinarily are not expressed on the same cell type |
| Mass cytometry | Unsupervised clustering and then used TREX to distinguish the immune markers that defined those clusters. Once identified, two-dimensional analysis are used to confirm. |
| Single-cell RNA sequencing | Cells identified by the doublet finder (A computational doublet detection method). We then used UMAPs and proximity to monocytes and B cells for example, to identify monocyte-T cell complexes and B cell-T cell complexes, transcriptionally. |

**Table S4. Percentage of cell types over total CD45^+^ immune cells stratified by metabolic disease**

|  | N | Non-Diabetics N=14 | Pre-diabetic and Diabetic N=24 | **Test** |
| --- | --- | --- | --- | --- |
| CD57^+^ CD161lo NK cells (1) | 38 | 7.5 [2.6, 10.7] | 5.9 [2.6, 10.8] | 0.7 |
| Classical monocytes (2) | 38 | **8.1 [6.6, 11.5]** | 6.1 [4.9, 7.1] | **0.02** |
| CD4 Naïve T cells (3) | 38 | 7.2 [5.5, 8.8] | 7.6 [2.2, 9.2] | 0.6 |
| CGC^+^ CD8^+^ T cells (4) | 38 | 6.0 [2.9, 7.9] | 6.1 [1.8, 9.7] | 0.7 |
| CD4 TEM/TH1 (5) | 38 | 5.3 [4.3, 7.9] | 4.6 [3.3, 6.4] | 0.2 |
| Mature B cells (6) | 38 | 5.3 [4.0, 6.5] | 3.8 [2.2, 6.6] | 0.07 |
| CGC^+^ CD4^+^ T cells (7) | 38 | 1.8 [0.6, 4.6] | **5.8 [1.7, 9.1]** | **0.007** |
| CD57^-^ CD161^+^ NK cells (8) | 38 | 3.2 [2.1, 3.6] | 3.7 [2.7, 7.1] | 0.31 |
| CD8^+^ Naïve (9) | 38 | **4.8 [3.4, 7.3]** | 3.5 [2.1, 4.3] | **0.02** |
| CD161^+^ CD4^+^ T cells (10) | 38 | 3.5 [2.7, 5.0] | 3.2 [1.8, 5.2] | 0.6 |
| CD8^+^ NKT cells (11) | 38 | 4.6 [1.3, 5.3] | 3.7 [0.9, 4.9] | 0.6 |
| CD8 TEMRA (12) | 38 | 4.9 [2.8, 5.5] | 3.1 [1.8, 4.5] | 0.2 |
| CD8 TEM (13) | 38 | 4.0 [3.1, 4.9] | 2.8 [1.4, 5.5] | 0.4 |
| CD8 TEMRA (14) | 38 | 3.4 [1.6, 5.0] | 3.2 [1.6, 5.0] | 0.7 |
| 𝛾𝛿 T cells (15) | 38 | **2.9 [2.0, 3.4]** | 1.5 [0.5, 2.3] | **0.01** |
| CD4 TEM (16) | 38 | 3.1 [2.3, 5.3] | 2.5 [1.7, 3.3] | 0.07 |
| CD57^-^ CD8^+^ TEM (17) | 38 | 2.7 [1.5, 3.3] | 1.6 [1.0, 2.6] | 0.2 |
| CD4^+^ T cell: CD14^+^ Monocyte complex (18) | 38 | 0.0050 [0.0025, 0.0069] | **0.014 [0.0075, 0.25]** | **0.003** |
| CD8^+^ T cell: CD14^+^ Monocyte complex (19) | 38 | 0.0063 [0.0025, 0.012] | **0.034 [0.0075, 0.22]** | **0.005** |
| NC Monocytes (20) | 38 | 1.9 [1.2, 2.5] | 1.9 [1.9, 2.7] | 0.6 |
| CD161^+^ CGC^+^ CD8^+^ T cells (21) | 38 | 1.0 [0.5, 3.3] | 1.4 [0.4, 3.0] | 1.0 |
| Memory B cells (22) | 38 | **2.1 [1.6, 2.4]** | 0.8 [0.6, 1.4] | **0.003** |
| CX3CR1^+^ GPR56^+^ CD57^-^CD4^+^ T cells (23) | 38 | 0.6 [0.4, 1.0] | **1.1 [0.7, 1.7]** | **<0.05** |
| CRTH2^+^ Intermediate monocytes (24) | 38 | 1.0 [0.5, 1.3] | 0.9 [0.5, 1.1] | 0.8 |
| CD4 T regulatory cells (25) | 38 | **1.4 [1.1, 2.2]** | 0.8 [0.5, 1.0] | **<0.001** |
| B cell; CD14^+^ Monocyte complexes (26) | 38 | 0.048 [0.025, 0.062] | **0.08 [0.04, 0.2]** | **0.03** |
| CD8^+^ T cell: CD14+ Monocyte complexes (27) | 38 | 0.0025 [0, 0.0025] | **0.009 [0.003, 0.09]** | **<0.001** |
| NK cell: Monocyte complexes (28) | 38 | 0.091 [0.054, 0.15] | 0.17 [0.10, 0.29] | 0.06 |
| CD3^+^ B cells (29) | 38 | 0.35 [0.29, 0.44] | 0.58 [0.34, 0.97] | 0.06 |
| CD161^+^ CD8^+^ NKT cells (30) | 38 | 0.21 [0.14, 0.31] | 0.40 [0.20, 0.69] | 0.1 |
| CD3^+^ CD4^-^ CD8^-^(31) | 38 | **0.51 [0.26, 0.66]** | 0.27 [0.16, 0.47] | **<0.05** |
| CD3^+^ T cell:CD14^+^ complexes (32) | 38 | 0.24 [0.096, 0.35] | 0.30 [0.15, 0.43] | 0.4 |
| CRTH2^+^ CD38^+^ NC Monocytes (33) | 38 | **0.33 [0.26, 0.39]** | 0.12 [0.092, 0.23] | **0.002** |
| Plasmablasts (34) | 38 | 0.076 [0.051, 0.099] | 0.12 [0.062, 0.16] | 0.2 |
| CD3^+^ CD14^+^ T cell: C Monocyte complexes (35) | 38 | **0.076 [0.060, 0.10]** | 0.041 [0.029, 0.059] | **<0.001** |
| PD1^+^CD8^+^ TCR𝛾𝛿 (36) | 38 | 0.025 [0.016, 0.041] | 0.020 [0.015, 0.028] | 0.2 |
| CD14^+^ CD16^+/-^ Monocytes (37) | 38 | **0.015 [0.013, 0.019]** | 0.0025 [0, 0.0075] | **<0.001** |
| Median [Lower and upper quartile] for continuous variables.  N is the number of non-missing values.  Tests used: Wilcoxon test  Abbreviations: NC non-classical, C classical | | | | |

**Table S5. Plasma cytokines by diabetes status**

|  | **N** | **Non-Diabetic N=14** | **Pre-diabetic/Diabetic N=24** | **Test Statistic** |
| --- | --- | --- | --- | --- |
| Interferon γ pg/ml | 29 | 4.3 [3.5, 5.7] | 4.6 [3.9, 6.2] | 0.59 |
| IL-12p70 pg/ml | 32 | 0.21 [0.16, 0.25] | 0.20 [0.17, 0.26] | 0.72 |
| IL-4 pg/ml | 27 | 0.05 [0.04, 0.06] | 0.05 [0.05, 0.06] | 0.77 |
| TNF-α pg/ml | 37 | 1.21 [0.89, 1.36] | 1.28 [0.94, 1.42] | 0.32 |
| IL-12/IL23p40 pg/ml | 29 | 100 [85, 115] | 105 [74, 140] | 0.66 |
| IL-17A pg/ml | 29 | 3.9 [3.3, 4.5] | 3.6 [2.9, 5.9] | 0.75 |
| IP-10 pg/ml | 19 | 113 [104, 133] | 102 [78, 128] | 0.69 |
| MIP-1α pg/ml | 29 | 12.7 [11.7, 13.4] | 11.5 [10.0, 15.1] | 0.86 |
| MIP-1β pg/ml | 29 | 64 [54, 86] | 63 [60, 81] | 0.79 |
| IL-1β pg/ml | 37 | 0.20 [0.16, 0.23] | 0.21 [0.16, 0.25] | 0.83 |
| IL-6 pg/ml | 37 | 1.47 [1.19, 1.92] | 1.44 [1.08, 1.79] | 0.69 |
| IL-10 pg/ml | 37 | 0.30 [0.27, 0.47] | 0.34 [0.26, 0.41] | 0.85 |
| Median [Lower and upper quartile] for continuous variables.  N is the number of non-missing values.  Tests used: Wilcoxon test | | | | |

**Table S6. Clonal TCRs and predicted antigens based on the closest matched CDR3 sequence**

| CDR3 | Total | Matched Sequence | Score | Epitope | Antigen | | | Source organism |
| --- | --- | --- | --- | --- | --- | --- | --- | --- |
| CASSDRDRDQPQHF | 8 | ASSVRDRDQPQH | 0.947 | GLCTLVAML^40^ | Transcriptional regulator IE63 homolog | | Human herpesvirus 4 (**EBV**) | |
| CAARLDTGGFKTIF | 5 | N/A |  |  |  | |  | |
| CSAKSPWTGGRNTEAFF | 4 | N/A |  |  |  | |  | |
| CASSFERQPQHF | 4 | ASSLSRQPQH | 0.936 | NLVPMVATV^41^ | HCMVUL83 | | Human herpesvirus 5 (**Human CMV**) | |
| CASSLLMASEQYF | 4 | ASSLLAADEQY | 0.9405 | GILGFVFTL^41^ | Matrix protein 1 | | **Influenza A virus** | |
| CASSLASGEQFF | 4 | ASSLASGEQY | 0.9907 | EHPTFTSQYRIQGKL^42^ | HCMVUL83 | | Human herpesvirus 5 strain Towne (**Human CMV**) | |
| CAFRSGSARQLTF | 4 | N/A | - | - | - | - | | |
| CVVSAYDYKLSF | 3 | N/A | - | - | - | - | | |
| CASKEGGAGGYTF | 3 | ASSEGRAGGYT | 0.9297 | FRDYVDRFYKTLRAEQASQE^43^ | Gag-Pol polyprotein | | Human immunodeficiency virus 1 (**HIV-1**) | |
| CARPPGGTGQDNYEQYF | 3 | NA | - | - | - | - | | |
| CASSQASGGADTQYF | 3 | ASSQARGGAETQY | 0.9377 | TSTLQEQIGW^44^ | gag protein | | Human immunodeficiency virus 1 (**HIV-1**) | |
| CASSLAGQSDGYTF | 3 | ASSLAGREGGYT | 0.9082 | SYFTNMFATWSPSKARLHLQ^45^ | Coagulation factor VIII | Homo sapiens (human) | | |
| CAVLDSNYQLIW | 3 | NA |  |  |  |  | | |
| CASGSDRGLAGVDTQYF | 3 | ASDQGLAGVDTQY | 0.9112 | FLNGSCGSV^46^ | orf1ab polyprotein [SARS CoV2] | SARS-CoV2 | | |
| CASGAGGNTEAFF | 3 | ASSQAGGNTEAF | 0.9358 | SYFTNMFATWSPSKARLHLQ^45^ | Coagulation factor VIII | Homo sapiens (human) | | |
| CASSLGQKNSPLHF | 3 | ASSLGQKNSPLH | 1.0000 | WPVTLACFV^47^ | Membrane protein | SARS-CoV2 | | |
| CSAKPGSEQFF | 3 | ASSPGTEQF | 0.9381 | IPSINVHHY^48^ | HCMVUL83 | Human herpesvirus 5 (**Human CMV**) | | |
| CASKAGGTYEQYF | 3 | ASMAGGTYEQY | 0.9625 | KLGGALQAK | 55 kDa immediate-early protein 1 | Human herpesvirus 5 strain AD169 (**Human CMV**) | | |
| CAASAPADNNNDMRF | 3 | NA |  |  |  |  | | |
| CASSLDRVNGYTF | 3 | ASSLDRGVGYT | 0.9120 | RAKFKQLL^49^ | Trans-activator protein BZLF1 | Human herpesvirus 4 (**EBV**) | | |
| CASGTIRAIYGYTF | 3 | ASSAIVAVYGYT | 0.9006 | KVCEFQFCNDPFLGVYYHKNNKSWMESEFRVYSSANNCTFEYV^46^ | SARS-CoV2 | SARSCoV2 | | |
| CASGKGSEQFF | 3 | ASGKGSEAF | 0.9735 | RQLLFVVEV^46^ | orf1ab polyprotein [SARSCoV 2] | SARS-CoV2 | | |
| CASSTPRTVYSNQPQHF | 3 | ASSPRTTYSNQPQH | 0.9223 | FVDGVPFVV^46^ | orf1ab polyprotein [SARSCoV2] | SARS-CoV2 | | |
| CASRQGARKTQYF | 3 | ASSRGASRETQY | 0.9044 | NLVPMVATV^41^ | HCMVUL83 | Human herpesvirus 5 (**Human CMV**) | | |

**Table S7. Clinical Demographics of HIV-negative Cohort #2**

| **Race** | **Gender** | **Age** |
| --- | --- | --- |
| White | Female | 52 |
| White | Female | 73 |
| White | Male | 68 |
| Black | Male | 69 |
| White | Male | 40 |
| White | Male | 78 |
| White | Male | 57 |
| White | Male | 65 |
| White | Male | 71 |
| Native Hawaiian or Other Pacific Islander | Male | 79 |

**Figure S1. Mass cytometry gating workflow.**

(A) Two-dimensional plots show the strategy used to gate PBMCs. After excluding control beads and selecting single cells, CD45 live cells were gated. Downstream of this, we performed UMAP analysis.
(B) Bi-axial plots of CD3^+^ T cell-CD14^+^ Monocyte complexes, CD3^+^ T cell-CD19^+^ B cell complexes, CD19^+^ B cell CD14^+^ monocyte complexes, and CD56^+^ NK cells-CD14^+^ monocyte complexes.
(C) Bi-axial plots show the gating strategy for CD4^+^ T cell helper subsets for TH1 (CXCR3), TH2 (CRTH2, CCR4) and TH17 (CD161, CCR6). Representative plots from non-diabetic and diabetic PWH are included.

**Figure S2. PBMCs from HIV-negative persons with diabetes detectable CD3^+^ T cell-CD14^+^ monocyte complexes**.

(A) (A)	Two-dimensional plots of flow cytometry data show the strategy used to gate PBMCs for cell sorting of CD3^+^ CD14^+^ complexes. After gating on cells of interest which include the lymphocyte and monocyte gates, live cells were gated. Downstream of this, we gated on CD3^+^ live cells, CD3^+^ T cells-CD14^+^ monocyte complexes, CD4^+^ and CD8^+^ T cells.

(B) Two-dimensional flow cytometry plots show CD3^+^ T cell-CD14^+^ monocyte complexes on three representative participants with HIV (non-diabetic (non-DM), pre-diabetic (PreDM) and Diabetic (DM) as a proportion of total live cells.

(C) Two-dimensional flow cytometry plots show CD3^+^ T cell-CD14^+^ monocyte complexes on three representative HIV-negative participants with diabetes as a proportion of total live cells.

(D) Violin plots show differences in CD3^+^ T cell-CD14^+^ monocyte complexes in PWH (Non-DM, PreDM/DM) and HIV-negative with diabetes.

Statistical analysis by Kruskall-Wallis test (C) and Mann Whitney test (D).

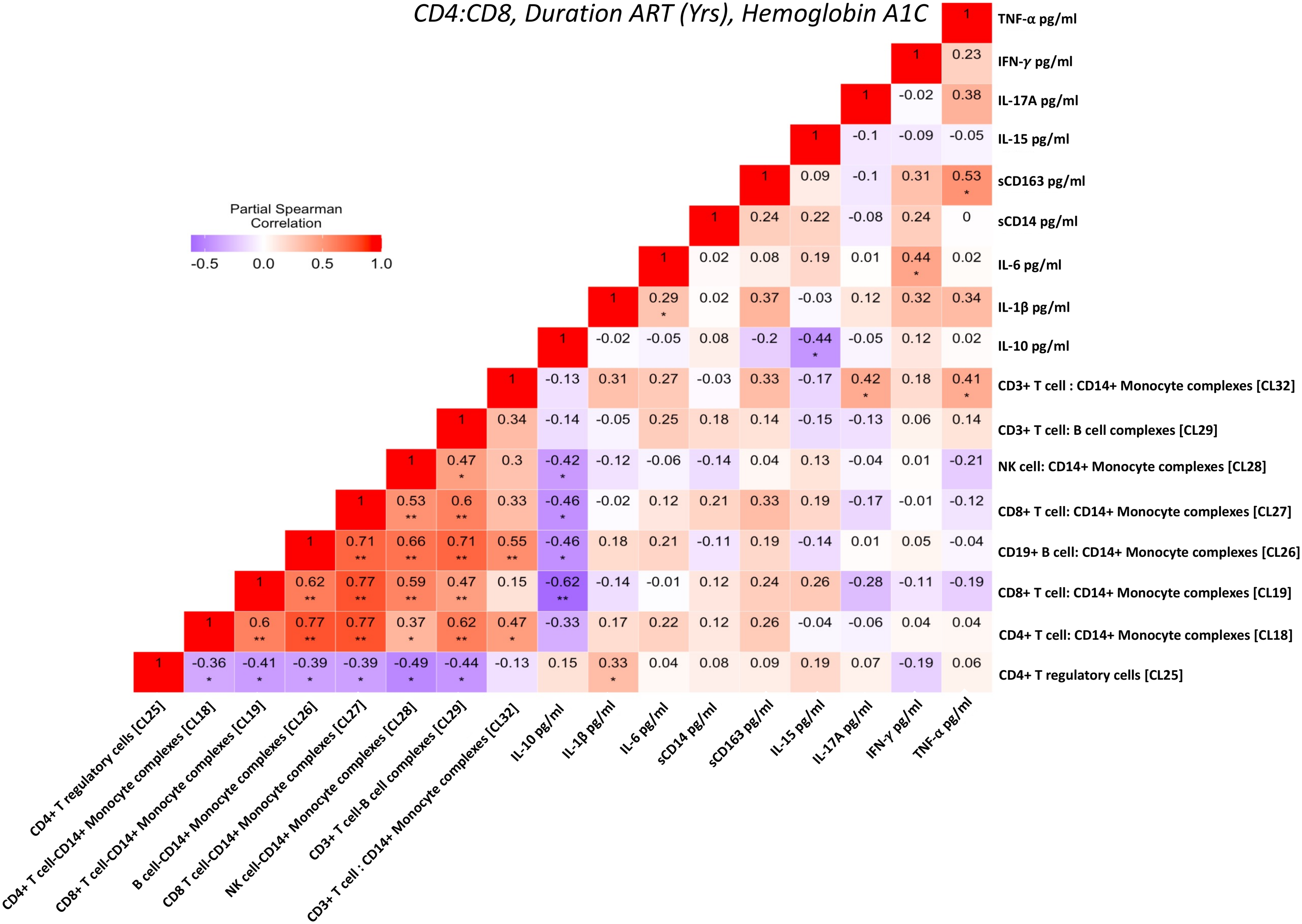

**Figure S3. The relationship between the cell-cell complexes and hemoglobin A1C is weaker for individuals with higher IL-10 or CD4^+^ T regulatory cells**.

The heatmap shows a partial Spearman correlation between cell-cell complex clusters, CD4^+^ T regulatory cells from 38 participants as defined by mass cytometry, and inflammatory cytokines (IL-10, IL-6, sCD14, sCD163, IL-15, IL-17, and IL-1β]) adjusted for CD4:CD8, antiretroviral therapy duration, and hemoglobin A1C (* p< 0.05, ** p<0.01).

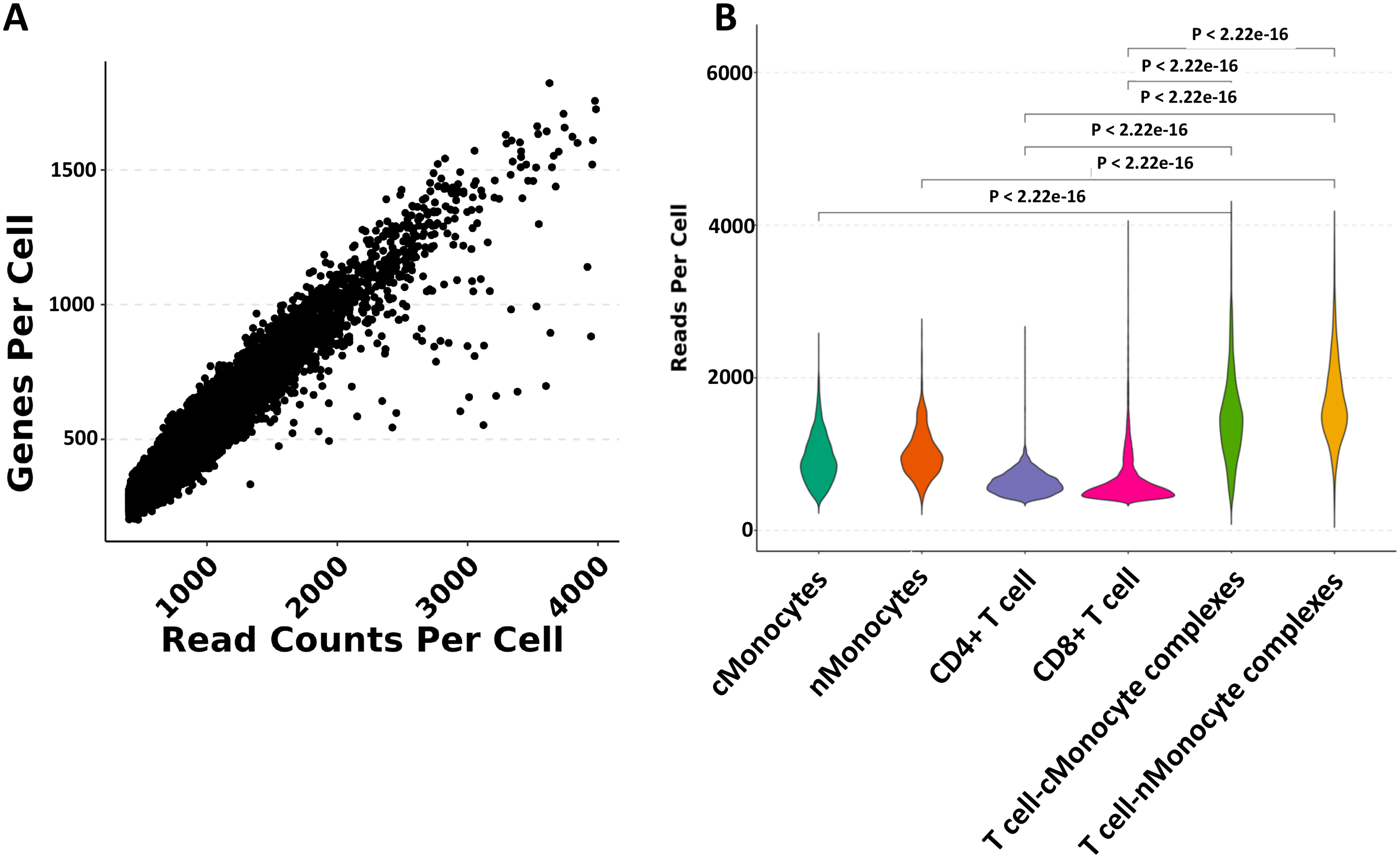

**Figure S4. T cell-monocyte complexes have more gene reads per cell than singlet T cells and monocytes**.

(A) Correlation plot between genes per cell and reads per cell count
(B) Violin plots show differences in gene reads per cell between singlet cell clusters and T cell-monocyte complexes in CITE-seq data.

Statistical analysis by Kruskal Wallis test (B).

**Figure S5. CD3^+^ T cells-CD14^+^ monocyte complexes are reduced after inhibition of oxidative phosphorylation**

(A) UMAP depicts protein translation by measuring puromycin uptake at baseline for each subset of cells [CD3^+^ CD14^+^, CD14^+^ monocytes, CD16^+^ monocytes, and CD4/CD8 T cell subsets (naïve, TCM, TEM, and TEMRA) in 15 individuals (blue: non-diabetic PLWH, magenta: 10 pre-diabetic, diabetic PWoH). The heatmap on the right shows the different markers that define each cluster.

(B) UMAP depicts the glucose dependence of each cluster, with a legend showing the differential scale.

(C) UMAP depicts the mitochondrial dependence of each cluster.

(D) Bar plots show percent glucose dependence (left) and percent mitochondrial dependence (right) of CD3^+^ T cells-CD14^+^ monocyte complexes, CD14^+^ Monocytes, CD4^+^ TCM, and CD8^+^ TCM (*blue dots – non-diabetic, magenta dots – Pre-diabetes/diabetes*).

(E) The proportion of CD3^+^ T cell-CD14^+^ monocyte complexes decreases with oligomycin's inhibition of oxidative phosphorylation. There is no change with 2DG.

Statistical analysis using Mann Whitney Test (D) and paired t-test (E).
